# Systems Genetics Cytokine Screen Identifies T cells as Necessary for Letrozole-Induced LH Elevation in a PMOS-like Mouse Model

**DOI:** 10.1101/2025.01.08.631835

**Authors:** Naveena Ujagar, Leandro Velez, Christy Nguyen, Kiara Wiggins, Gabriela De Robles, Zena Del Mundo, Siaje Gideon, Jessica Cassin, Joshua Kim, Nandini Naidu, Julio Ayala Angulo, Benjamin Babaev, Alexius Dingle, Margareta D. Pisarska, Ricardo Azziz, Jessica L. Chan, Rachel A. Ross, Alexander S. Kauffman, Varykina G. Thackray, Beata Banaszewska, Ewa Wysocka, Antoni Duleba, Marcus Seldin, Dequina Nicholas

## Abstract

Polycystic ovary syndrome (PMOS) is a complex reproductive disorder with clear genetic susceptibilities that impact the heterogeneous clinical presentation of symptoms and severity through unknown mechanisms. Chronic inflammation is linked to PMOS, but a clear cause-and-effect relationship between immune mediators and PMOS phenotypes has yet to be demonstrated. This study employed a comprehensive systems immunology approach, utilizing a letrozole-induced PMOS mouse model to identify changes in inflammatory factors associated with PMOS symptoms. By analyzing immune cells and secreted cytokines from 22 different mouse strains, we identified T cells and TNF-β as associated with PMOS-like phenotypes, regardless of genetic background. We used a knockout of TCRα to show that functional T cells are necessary for development of pathologically elevated luteinizing hormone (LH) in letrozole-treated female mice. In women with PMOS, we observed elevated TNF-β transcripts in immune cells from women with PMOS. Finally, we demonstrate that TNF-β increased Lhb mRNA in a female mouse gonadotrope-derived cell line, suggesting that TNF-β may directly modulate gonadotrope gene expression and may contribute to elevated LH in PMOS-like conditions. These findings support a requirement for functional αβ T cells in LET-induced LH elevation in a PMOS-like mouse model and identify TNF-β as a candidate immune mediator for further investigation.

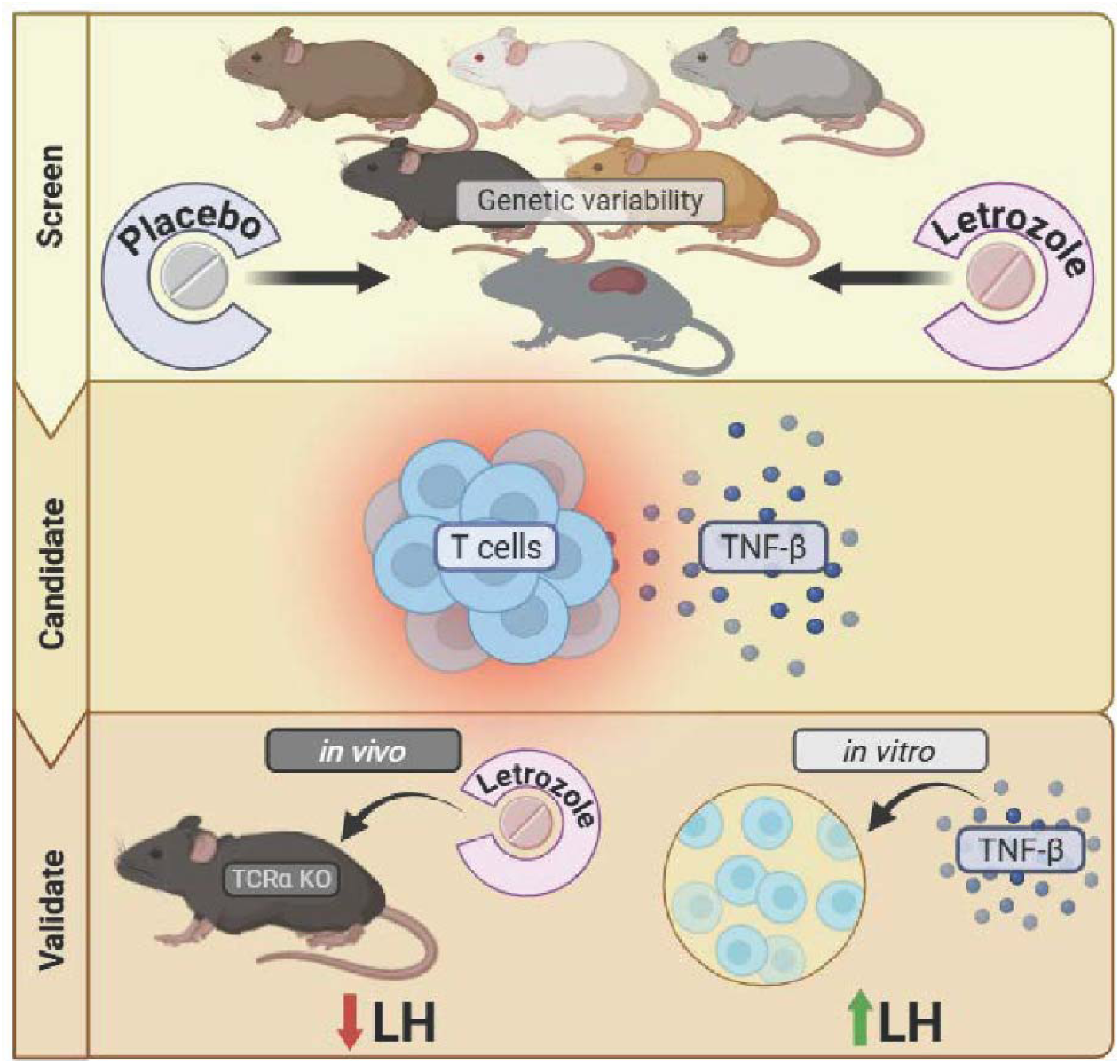

**One Sentence Summary:** Functional αβ T cells are linked to LET-induced LH elevation in a PMOS-like mouse model, uncovering candidate immune mechanisms for further study.

## INTRODUCTION

Polyendocrine Metabolic Ovary Syndrome (PMOS, formerly known as Polycystic Ovary Syndrome, or PCOS) is the most common reproductive disorder in women of childbearing age, affecting as many as 10-15% of women (*1*). PMOS is characterized by polycystic ovarian morphology (PCOM), oligo- or anovulation, and clinical or biochemical hyperandrogenism, which ultimately results in subfertility (*2*). Clinically, women with PMOS present heterogeneously, and PMOS is categorized as types A-D based on three primary features: hyperandrogenism, irregular cycles (an- or oligo-ovulation), and PCOM (*2–5*). Type A or Classic PMOS is the most prevalent and most studied subtype of PMOS and is characterized by presentation of all three primary features and is often accompanied with symptoms like hirsutism, acne, reduced fertility, and metabolic syndrome. Women with Type B or non-PCOM PMOS have hyperandrogenism and irregular cycles but no PCOM. Type C or Ovulatory PMOS presents with hyperandrogenism and PCOM but regular ovulation. Finally, Type D or non-androgenic PMOS is characterized by ovulatory dysfunction and PCOM without elevated levels of androgen.

PMOS patients also exhibit elevated C reactive protein (CRP), a broad indicator of inflammation, which is 96% higher in women with PMOS than those without (*6*). Currently, CRP is the most reliable marker of chronic inflammation in PMOS (*6, 7*). Across studies, CRP is consistently upregulated in women with PMOS (*6–8*). However, the literature contains conflicting findings regarding the mediators of this chronic inflammation (*9*). Despite consistency in elevated CRP, no differences in circulating cytokines, such as IL-6, were detected in women with and without PMOS in several cohorts, though this was likely a consequence of small sample size (*10–12*). However, a meta-analysis of BMI-matched cohorts revealed significantly increased IL-6 in women with PMOS that correlated with insulin resistance (*13*).

The latter outcome is supported by a report demonstrating that administration of exogenous androgen to healthy women to an elevated level observed in PMOS increases circulating IL-6 (*14*). Clinical studies have begun to define inflammation in PMOS by identifying increased circulating levels of IL-1α, IL-1β, TSP-1 and TGF-β1 in PMOS (*15, 16*). Additionally, circulating neopterin (a protein made by macrophages) is elevated in PMOS independent of BMI (*17*). Collectively, the clinical literature provides emerging evidence of changes in circulating cytokines in PMOS but fails to inform on whether they contribute to PMOS symptomology, and which cells are involved, questions better addressed using a pre-clinical animal model.

Kauffman et al. developed a mouse model that utilizes chronic treatment with letrozole (LET), a nonsteroidal aromatase inhibitor, initiated in peripubertal life to induce the reproductive and metabolic hallmarks of Type A PMOS in female C57BL/6 mice (*18*). Aromatase inhibition by letrozole prevents the conversion of androgens to estrogens, thereby resulting in elevated circulating androgen levels that mimic the hyperandrogenism observed in PMOS patients. In addition to eliciting high androgen levels, the LET PMOS-like mouse model is particularly effective in inducing other critical features of the disorder, including irregular estrous cycles, polycystic ovaries, anovulation, and metabolic dysfunction such as insulin resistance and body weight gain (*18, 19*). The LET model exhibits at least two characteristics that are comparable to the Rotterdam criteria used to diagnose PMOS in women (*20*). This model also successfully recapitulates underlying endocrine mechanisms observed in women with PMOS, such as hyperactive luteinizing hormone (LH) pulsatility (*21, 22*).

Recent studies have highlighted the complex interplay between genetic predisposition and environmental factors in shaping the diversity of PMOS phenotypes. For instance, a genome-wide association study by Day et al. identified 14 novel loci associated with PMOS, suggesting a strong genetic component to the syndrome (*23*). However, the expression of these genetic variants can be significantly influenced by environmental factors. Lifestyle factors such as diet and exercise have been shown to modulate PMOS symptoms. For example, weight loss through lifestyle intervention can improve metabolic and reproductive outcomes in women with PMOS, regardless of their genetic predisposition, implicating conserved mechanisms (*24*). Furthermore, epigenetic modifications, which can be influenced by both genetic and environmental factors, have been implicated in PMOS pathogenesis. Kokosar et al. identified altered DNA methylation patterns in adipose tissue of women with PMOS, suggesting that epigenetic changes may contribute to the metabolic dysfunction often observed in the syndrome (*25*). These findings underscore the importance of considering both genetic and environmental factors when studying PMOS phenotypes to decipher mechanisms specific to disease development.

The high variability and severity of PMOS phenotypes across women with the disorder poses challenges in identifying immune-related mechanisms specific to PMOS. Using the LET-induced model of PMOS and a genetic screen, we aimed to address the challenge of identifying immune mediators specific to PMOS and the relationship of those immune mediators to diverse PMOS phenotypes. Here, we perform a comprehensive and robust analysis of immune cells and cytokines associated with PMOS symptoms. Using a systems genetics approach by analyzing placebo- and LET- treated female mice across 22 distinct inbred mouse strains, we 1) identify that functional αβ T cells are required for LET-induced LH elevation in a PMOS-like mouse model and 2) uncover TNF-β as a candidate immune modulator of LH secretion in PMOS-like conditions.

## RESULTS

### Confirmation of PMOS-like phenotypes in LET-induced mouse model in multiple strains

To discover immune regulators in PMOS, we sought to perform correlations and regression analyses of immune profiles with PMOS-like symptoms in mice. However, traditional approaches using single or few strains would be under-powered for these types of analyses given the lack of variation within inbred strains. To address this limitation and enhance statistical power, we used the established approach of chronic LET treatment across 22 inbred strains selected for their genetic and immune diversity (*26–28*). The strains selected have demonstrated variance in their metabolic response to high fat diet or have differing immune cell composition compared to the commonly used C57BL/6 mouse. This multi-strain design creates sufficient variability for correlation analyses, though it remains underpowered to detect within strain differences. It is important to note that the small sample sizes within individual strains (ranging from 2 to 4 mice per group, as detailed in Supplemental Table 1) preclude within-strain comparisons. The statistical power of this study is specifically designed for cross-strain correlation analyses. We implanted LET or placebo (control) pellets in 130 young adult female mice at ∼9 to 10 weeks of age and replaced with a fresh pellet after 3 weeks to ensure continuous delivery through the entire 6-week paradigm (mouse cohort 1) (**Fig. 1A**). The mice were phenotypically characterized during the final two weeks of treatment and immune profiles for each animal were generated by immunophenotyping inguinal lymph nodes and performing cytokine profiling on immune stimulated spleen cultures (**Fig. 1A**). Across the entire cohort, LET treatment *globally* induced PMOS-like symptoms in these strains. As expected, compared to the placebo controls, LET significantly increased body weight (BW) gain over time, with a slight temporary plateau due to disruption in LET release kinetics following the week 3 re-implantation surgery (**Fig. 1B**). LET mice on average gained 2-fold more BW than placebo control females and had significantly larger inguinal fat pads (**Fig. 1C and D**), confirming prior reports (*18, 19, 29*). This increase in BW was accompanied by impaired glucose tolerance, a common metabolic symptom of PMOS (**Fig. 1E and 1F).** Blood serum testosterone, of which elevated levels are often a primary diagnostic criterion for PMOS, were also significantly increased in LET mice compared to controls, recapitulating hyperandrogenism in PMOS women (**Fig. 1G**). LH was significantly increased as previously reported (*18*) (**Fig. 1H**). The LH to FSH ratio, though not statistically significant, displayed a trend towards an increase in LET mice (p=0.063), as occurs in many women with PMOS (**Fig. 1I**). Finally, the ovaries of LET mice had a significant increase in cystic follicles and fewer corpora lutea (an indicator of ovulation), consistent with ovarian morphology and oligo/anovulation in women with PMOS (**Fig. 1J, 1K**). Compared to published outcomes in LET treated C57BL/6 mice (*18, 29*), our data demonstrate significant variability across PMOS-like symptoms, indicative of phenotypic diversity sufficient to power correlation analyses.

**Figure 1.**
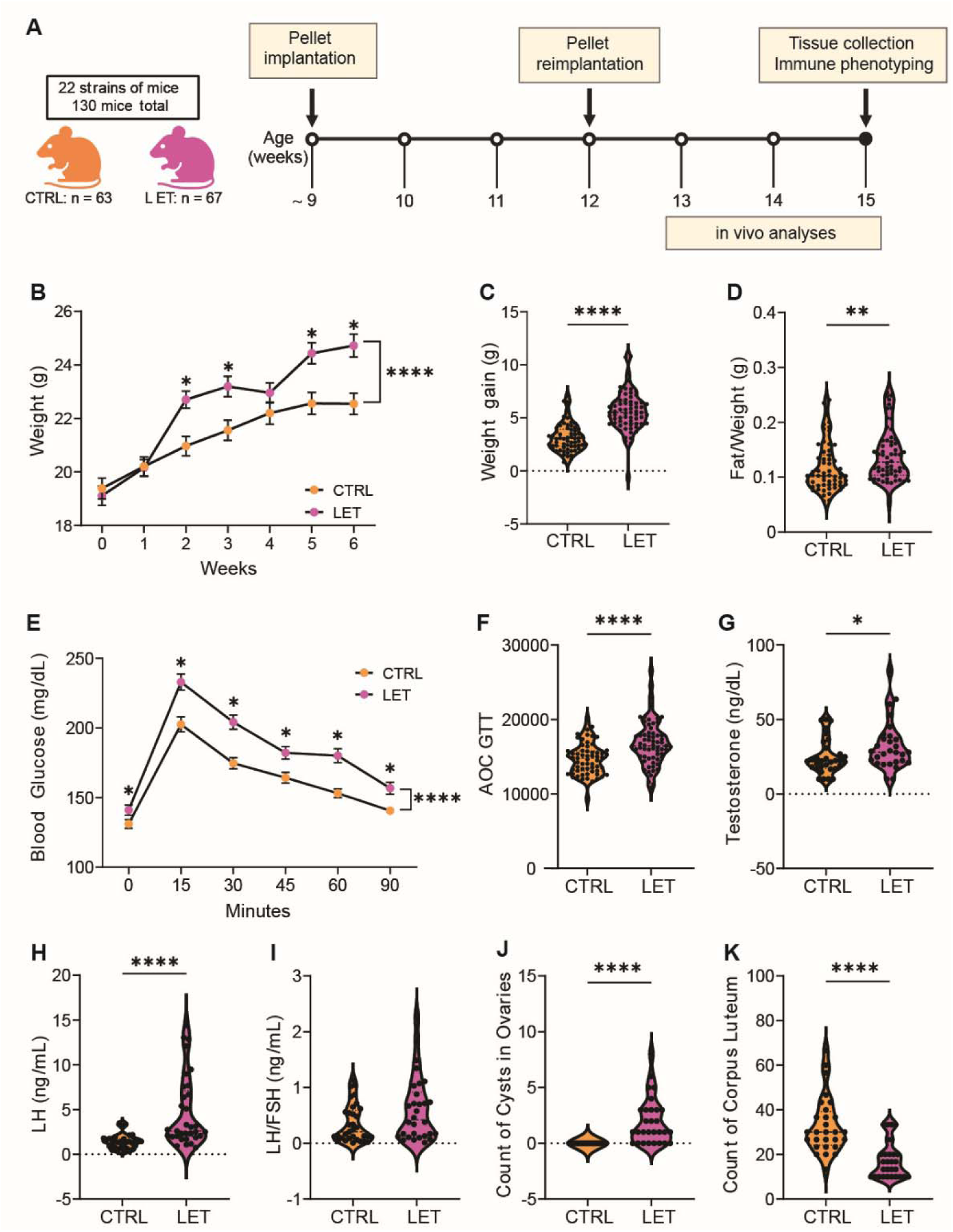
Confirmation of PMOS-like symptoms in LET-induced mouse model in multiple strains. (**A**) Timeline of cohort 1 letrozole or placebo pellet implantation at week 9 and 12. *In vivo* analyses (glucose tolerance testing, body composition, and estrous cycling) were measured between 13 to 15 weeks, and sac at 15 weeks included collection of spleen, inguinal lymph nodes and serum. (**B**) Total body weight during the 6 weeks of treatment. (CTRL=63, LET=67). (**C**) Overall weight gain over a 6-week period (CTRL=63, LET=67). (**D**) Fat to total body weight ratio (CTRL=63, LET=67). (**E**) Blood glucose (mg/dL) in response to a glucose tolerance test (CTRL=61, LET=65). (**F-I)** Serum hormone measurements: (**F**) Area under the curve of glucose tolerance testing (CTRL=63, LET=67). (G) Testosterone (ng/dL) (CTRL=32, LET=31). (**H**) Luteinizing hormone (ng/mL) (CTRL=33, LET=30). (**I**) Ratio of luteinizing hormone to follicular-stimulating hormone (ng/mL) (CTRL=33, LET=30). (**J**) Count of cystic follicles per ovary (CTRL=27, LET=29). (**K**) Count of corpora lutea per ovary (CTRL=28, LET=29). Data are presented as mean ± SEM and analyzed with t-test for parametric data or Mann-Whitney test for nonparametric data. Significance was accepted at p<0.05 and is indicated with an *.

### Immune and cytokine profiles implicate T cells in development of PMOS phenotypes

The large genetic diversity of this study provides variation and robust statistical power to identify novel relationships between immune variables and phenotypic outcomes. We performed immunophenotyping of mouse spleen and inguinal lymph nodes in LET and control mice (**Fig. 1A**). Cells from spleen and lymph nodes from mouse cohort 1 were stained with a 18-color antibody panel and analyzed by flow cytometry (**Supplemental Fig. 1 and 2**). The global distribution of immune cell populations in inguinal lymph nodes or spleen did not shift with LET-induction of PMOS-like phenotypes in the diverse cohort 1 (**Supplemental Fig. 2 A-B**). Because genetic strain alone substantially impacts the basal immune cell distribution of immune cells in the spleen (*30*) and because the study is not powered to assess immune frequencies by strain, we performed principal component analysis (PCA) of immune frequencies and confirmed that variability within the dataset was driven by strain (**Supplemental Fig. 2 C-D**). These results are expected and consistent with established literature on inbred mouse strain immune variability (*31–33*). These well-characterized strain differences in immune composition were intentionally leveraged to generate the phenotypic diversity necessary to power cross-strain correlation analyses. Since this systems genetics approach is intentionally designed for cross-strain correlation analyses and is not powered for within-strain comparisons, we correlated lymph node immune cell frequencies with PMOS-like traits in control or LET mice. Notably, the association between memory CD4+ T cells and testosterone was observed in both control and LET-treated animals, suggesting this relationship reflects a broader immunoendocrine interaction rather than a PMOS-specific phenomenon. The opposite directionality of this correlation between control and LET mice raises the possibility that this immunoendocrine relationship may be altered under PMOS-like conditions, though this interpretation warrants further investigation (**Fig. 2A**). In control animals of cohort 1, memory CD4+T cells positively correlated with testosterone, but this correlation flipped to negative in LET-treated mice, implying a change in T cell function in PMOS-like conditions.

**Figure 2.**
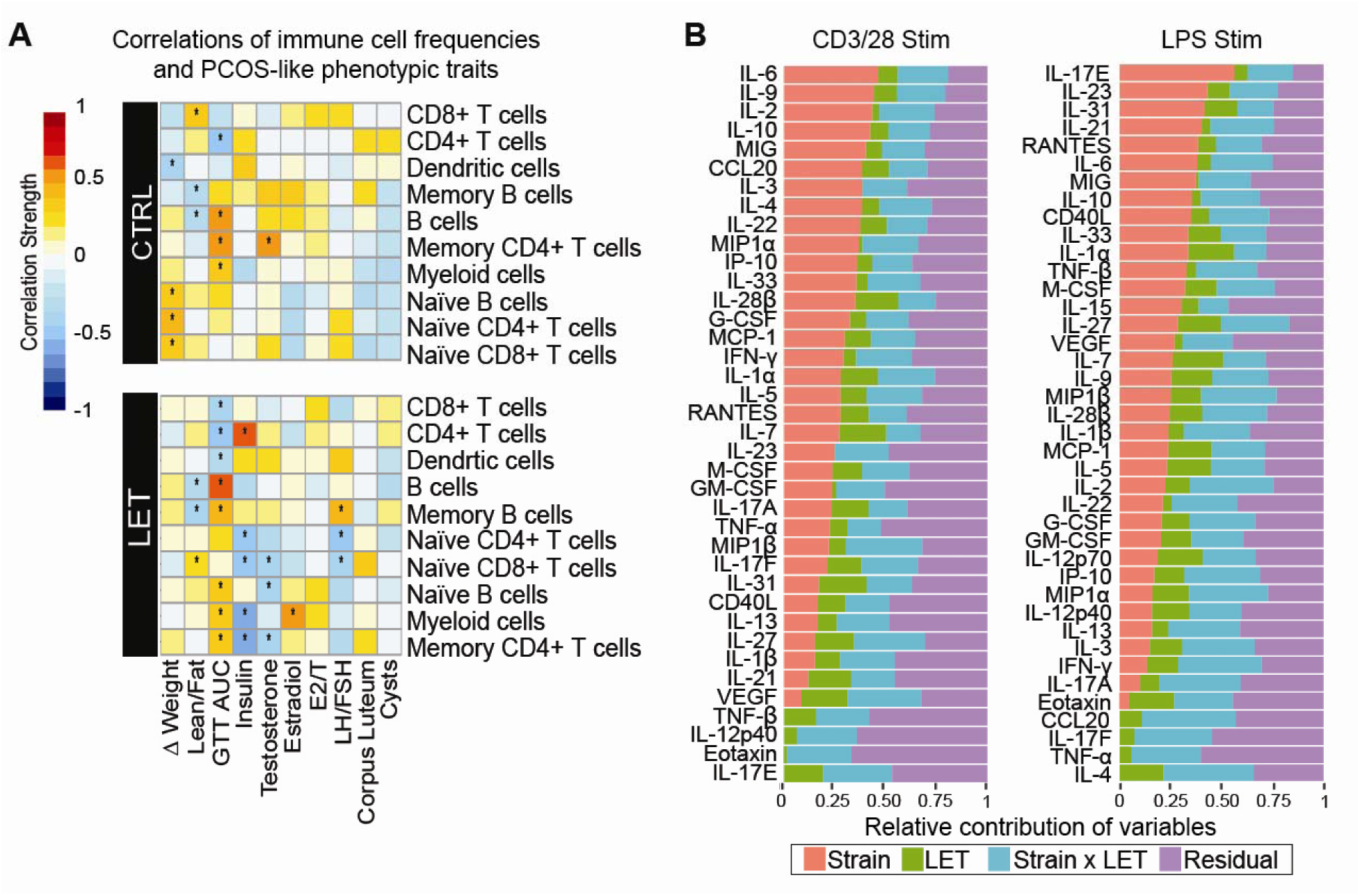
Immune and cytokine profiles implicate T cells in development of PMOS phenotypes. (**A**) Heatmap depicting correlative relationships between flow cytometry enumerated lymph node immune cell frequencies and PMOS-like phenotypes from the genetically diverse CTRL and LET cohort 1. Red denotes strong, positive correlation. White denotes no correlation. Blue denotes strong, negative correlation. Asterisk (*) denotes statistically significant correlations. (**B**) Heritability estimates across the genetically diverse cohort of cytokine concentrations from the conditioned media of splenocytes stimulated with anti-CD3/28 (Left) or LPS (Right).

We used similar approaches to analyze cytokine profiles generated *in vitro*. We cultured dissociated spleen in the presence of anti-CD3/CD28 (T cell stimuli) or Lipopolysaccharide (LPS, myeloid cell stimuli which targets Toll Like Receptor 4, TLR4) and analyzed the conditioned media for 40 secreted cytokines. Partial least squares discriminant analysis (PLSDA) showed that within many strains, the placebo and LET mice clustered together. In contrast, several strains displayed divergent cytokine profiles upon LET treatment in cohort 1 (**Supplemental Fig. 3**). For example, in LPS stimulated splenocytes, BXD73 and NOD strains had cytokine profiles that result in their placebo data point being located far from the centrality of the cluster for controls. Further, cytokine profiles did not shift for BALB/cj mice in response to LET. However, LPS-induced cytokine profiles from DBA mice displayed a clear shift. This strain variability in cytokine response to LET treatment powered multiple correlation analyses.

We tested whether reproductive and metabolic outcomes associated with PMOS correlated with cytokines measured from T cell or LPS stimulated immune cells from the mice in cohort 1(**Supplemental Fig. 4**). In control mice, MCP-1 and other inflammatory cytokines known to be associated with ovarian tissue remodeling were positively correlated with follicular cysts. This relationship, likely indicative of homeostatic mechanisms, was not present in PMOS-like mice. Interestingly, we observed that a subset of T cell cytokines (IL-33, IL-17E, CD40LG, IL-27, and TNF-β) were positively correlated with lean/fat ratio in either T cell or myeloid stimulation conditions. However, we did not identify any conserved cytokine signatures that strongly associated with reproductive or metabolic PMOS-like outcomes. Therefore, we took another approach to identify potential cytokines related to PMOS-like phenotypes. We performed a formal heritability estimate analysis using a linear mixed effects model across all of the strains and two treatment groups of mouse cohort 1 to determine which cytokine signatures are explained by genetics, LET treatment, or both (**Fig. 2B**). Heritability estimates for cytokines across all strains determined that some cytokines were strain independent, with variability only explained by LET treatment or the interaction of LET with the genetic background (**Fig. 2B**).

These cytokines included TNF-β, IL-12p40, Eotaxin, and IL-17F from T cell stimulated conditions and CCL20, IL17-F, TNF-α, and IL-4 from LPS stimulated conditions. Together, these findings prioritize T cells as candidate immune contributors to PMOS-like endocrine phenotypes, based on correlation between memory CD4+ T cells and testosterone and the predominance of T cell-derived cytokines among strain-independent contributors.

### Functional **αβ** T cells are required for LET-induced LH elevation and impaired estrous cyclicity

Given our data and a prior report that adoptive transfer of B cells from women with PMOS into mice and knockout of B cells in a PMOS-like model did not alter disease (*34*), we focused on testing the contribution of T cells to the development of PMOS-like phenotypes by utilizing the LET paradigm in young adult T cell receptor alpha (TCRα) KO mice using the same paradigm as in Figure 1 (mouse cohort 2) (**Fig. 3**). TCRα KO mice generate T cells but do not have a functional αβ T cell receptor and therefore lack αβ T cells. Outcomes from the present TCRαKO cohort were compared to C57BL/6 WT controls including mice from our previously published cohort of adult C57BL/6 mice (*29*). Female WT C57BL/6 mice treated with chronic LET in adulthood as opposed to peri-pubertally develop a weak metabolic phenotype but still exhibit a strong PMOS-like reproductive phenotype (*29*). Like adult WT C57BL/6 mice given LET, cohort 2 TCRα KO mice treated with LET in adulthood did not have a notable metabolic phenotype compared to placebo, as expected (**Supplemental Fig. 5A-C**). However, in contrast to LET WT females, LET TCRα KO females had ameliorated reproductive phenotypes (**Fig. 3**). As we and others have previously published, LET WT female mice do not cycle through estrous stages and remain arrested in diestrus (*18, 19, 29*) (**Supplemental Fig. 5D-E**), mimicking menstrual cycle disruption in women with PMOS. In contrast, cohort 2 LET TCRα KO females were not arrested in diestrus and had intact estrous cycles compared to Placebo females (**Fig. 3A-B**). The lack of T cells in TCRα KO females did not alter LET-induced elevation of testosterone (**Fig. 3C)** but was protective for the pathologically hyperactive LH secretion observed in LET-treated female mice similar to women with PMOS (**Fig. 3D**). FSH in TCRα KO females was lower than wild type controls but did not differ by LET treatment as expected (**Fig. 3E**). LET treated WT C57BL/6 females typically exhibit a complete loss of corpora lutea, increased cystic follicles, and hemorrhagic cysts compared to control (*19, 29*) as replicated by WT C57BL/6 sentinel mice (**Fig. 3F**). In contrast, some LET treated TCRα KO retained corpora lutea even in the presence of increased cystic follicles and hemorrhagic cysts (**Fig. 3G-J**). Taken together, data from the TCRα KO mice suggest that functional αβ T cells are necessary for some PMOS-like reproductive phenotypes, particularly elevated LH and impaired estrous cyclicity in the LET PMOS model.

**Figure 3.**
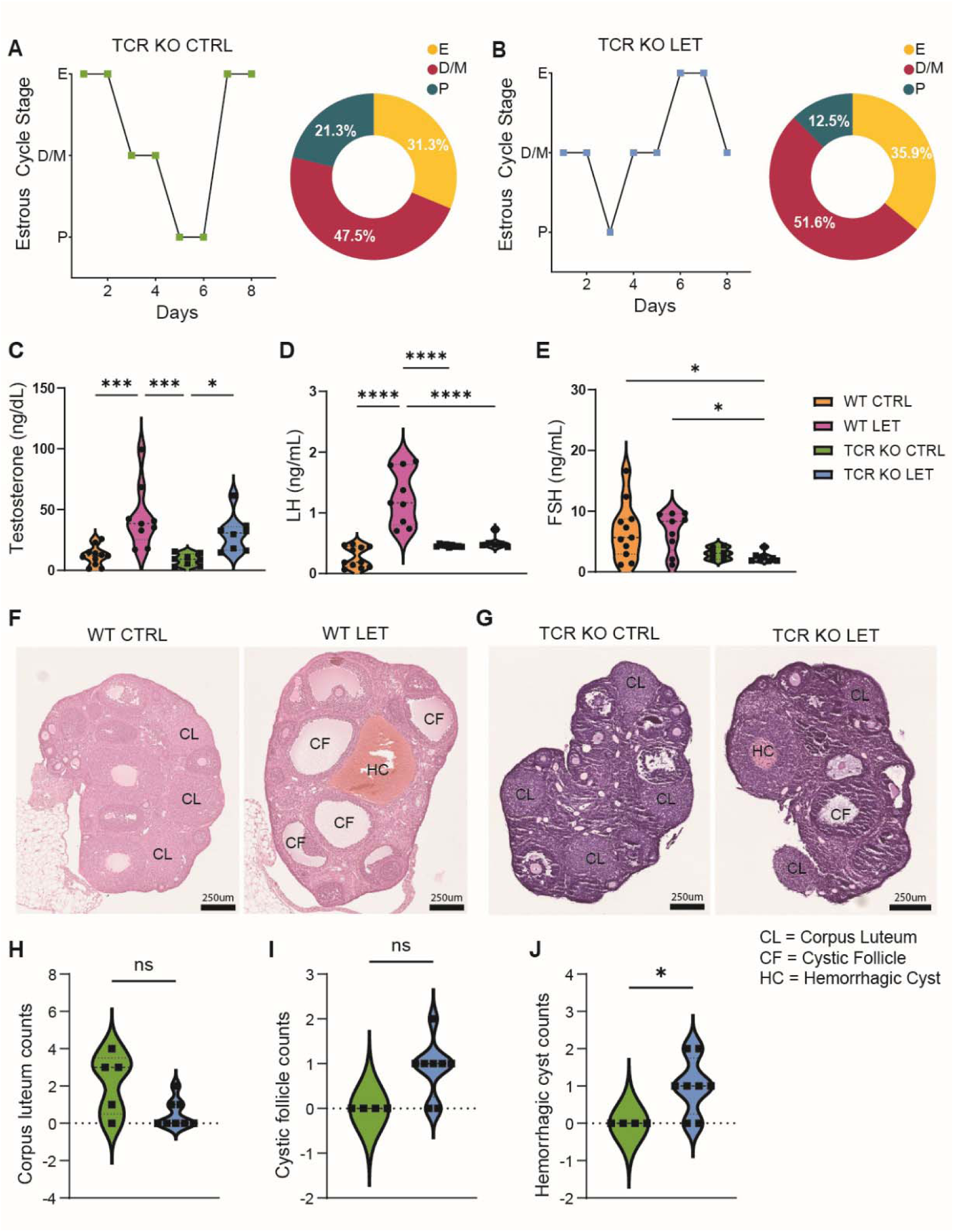
T cell KO prevents LET induced elevation of LH. (**A-B**) Representative estrous cycle plots and percent of mice cycling, defined as progressing through all estrous cycle stages in order of TCR KO CTRL (A) and TCR KO LET (B) within cohort 2. (**C-E**) Serum levels of testosterone (ng/dL) (C) luteinizing hormone (ng/mL) (D) and follicular-stimulating hormone (ng/mL) (E). Data analyzed with 2-way Anova. Significance was accepted at p<0.05 and is indicated with an *. (**F-G**) Representative images of ovaries from WT mice (F) and TCR KO mice (G). (**H-J**) Counts of ovarian features in TCR KO mice, including corpus luteum (H), cystic follicles (I) and hemorrhagic cysts (J). Data analyzed with t-test for parametric data or Mann-Whitney test for nonparametric data. Significance was accepted at p<0.05 and is indicated with an *.

### TNF-**β** is linked to adiposity and modifies androgen-associated clinical features in PMOS

Given that TCRα KO partially protects from abnormally elevated LH, we sought to identify potential cytokine mediators. We narrowed the list of cytokines identified by correlation and heritability analysis through the genetic screen to T cell-derived cytokines and tested whether these cytokines were increased in women with PMOS. Peripheral blood mononuclear cells (PBMCs) from a cohort of adult women with (n=15) and without (n=5) Type A PMOS were analyzed (human cohort 1). As expected, women with PMOS had elevated testosterone, fewer menses per year and larger ovarian volume compared to controls (**Table 1, Supplemental Fig. 6)**. This cohort of women with PMOS did not have a strong metabolic phenotype, similar to the adult murine model of LET-induced PMOS (*29*) (**Fig. 4**), and there were no significant group differences in insulin response or GTT (**Table 1**, **Fig. 4B-D**), though the lipid profiles of these women with PMOS were characterized by higher triglyceride levels and lower HDL cholesterol (**Fig. 4E-H**). We performed qPCR for TNF-β, IL-17E, IL-9, IL-27, CD40LG, IL-33, and IL-1α on the mRNA extracted from PBMCs given that T cells can secrete these cytokines, though T cells may not be the sole or primary source (*35*). TNF-β mRNA (*LTA*), one of the three cytokine transcripts that were detectable, was significantly increased in this small cohort of women with PMOS (**Fig. 4I**), consistent with a recent finding that TNF-β is elevated only in serum of lean women with PMOS (*36*).

**Figure 4.**
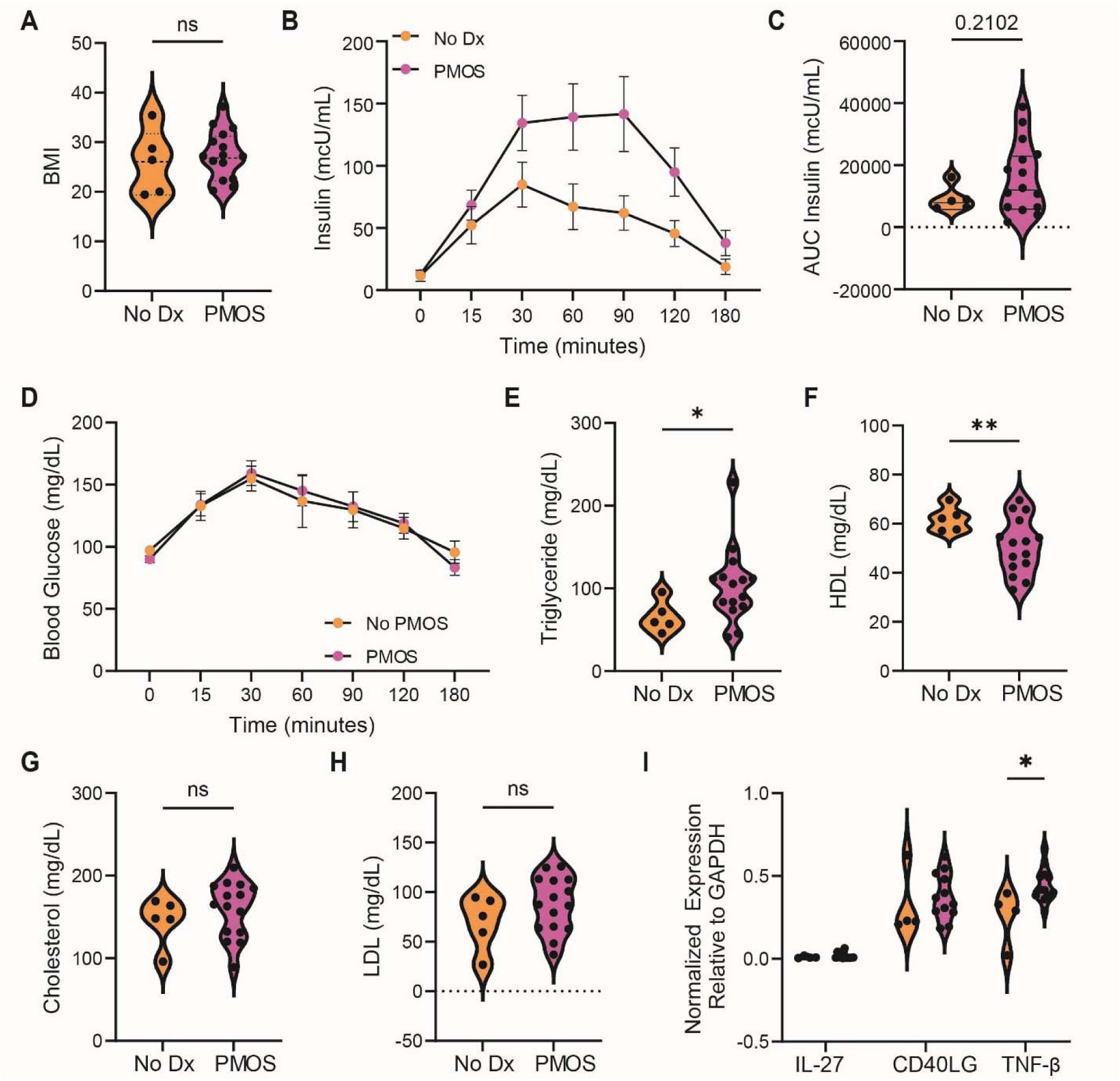
TNF-β mRNA expression is higher in PBMCs of women with PMOS. (**A-H**) Clinical parameters of the PMOS cohort. (A) Body mass index, BMI, (B) Insulin response, (C) Area under the curve of insulin response from (B), (D) Glucose tolerance test, (E) total triglycerides, (F) total high density lipoproteins, (G) total cholesterol, and (H) total low density lipoproteins measured in women with no PMOS diagnosis (n = 5) and with PMOS diagnosis (n = 15). (**I**) qPCR of mRNA expression of pro-inflammatory cytokines (IL-27, CD40LG, TNF-β) measured in PBMCs of women with no PMOS diagnosis (n = 4) and with PMOS diagnosis (n = 12). Data are presented as mean ± SEM and analyzed with t-test for parametric data or Mann-Whitney test for nonparametric data. Significance was accepted at p<0.05 and is indicated with an *.

**Table 1.** PMOS associated clinical parameters of women with no PMOS diagnosis (n = 5), or with PMOS diagnosis (n = 15). Data are presented as mean ± SEM and analyzed with t-test for parametric data or Mann-Whitney test for nonparametric data. Significance was accepted at p<0.05 and is indicated with an *.

|  | No PMOS Dx | PMOS | P-value | Significant? |
| --- | --- | --- | --- | --- |
| Age | 30.4 ± 3.91 | 25.93 ± 3.69 | 0.0604 | NS |
| BMI | 25.64 ± 6.64 | 27.25 ± 4.91 | 0.4973 | NS |
| Menses/year | 12 ± 0 | 7.07 ± 2.52 | <0.0001 | **** |
| R ovarian volume | 6.88 ± 3.12 | 14.19 ± 4.48 | 0.0035 | ** |
| L ovarian volume | 7.96 ± 3.15 | 13.66 ± 6.37 | 0.0328 | * |
| Testosterone | 0.32 ± 0.10 | 0.75 ± 0.26 | 0.0003 | *** |
| SHBG | 66.16 ± 30.30 | 43.55 ± 25.89 | 0.1186 | NS |
| DHEAS | 4.59 ± 1.68 | 9.70 ± 4.91 | 0.0025 | ** |
| LH | 7.44 ± 2.54 | 11.46 ± 6.78 | 0.3486 | NS |
| FSH | 6.72 ± 2.18 | 5.84 ± 1.99 | 0.5531 | NS |
| E2 | 57.84 ± 15.45 | 60.75 ± 24.22 | 0.866 | NS |
| PRL | 21.01 ± 4.81 | 20.99 ± 5.59 | 0.9466 | NS |
| Cholesterol | 139.86 ± 39.86 | 160.94 ± 29.53 | 0.3486 | NS |
| Triglycerides | 83.60 ± 36.91 | 97.95 ± 45.53 | 0.0178 | * |
| HDL | 50.70 ± 12.50 | 54.75 ± 10.95 | 0.0093 | ** |
| LDL | 72.46 ± 35.33 | 86.64 ± 25.81 | 0.2504 | NS |
| TSH | 1.79 ± 0.45 | 1.23 ± 0.60 | 0.0003 | *** |

Because we cannot determine which cell types are expressing *LTA* mRNA from bulk PBMCs, we used data from the Genotype-Tissue Expression (GTEx) project to infer likely cell types involved in TNF-β production in PMOS (*37*). GTEx is a well-powered transcriptomic research initiative aimed at understanding the relationship between genetic variation and gene expression across different human tissues. GTEx transcriptomic data of premenopausal women (n=31, age <50), demonstrated that *LTA* is highly expressed in both B and T cells. In particular, alpha beta memory CD4 T cells had extremely high expression (**Supplemental Fig. 7A**). Spleen *LTA* expression correlated with gene expression across multiple tissues, with visceral adipose tissue showing the greatest number of significant associations (**Supplemental Fig. 7B**).

The high number of adipose genes co-correlated with spleen expression of *LTA* indicated that TNF-β expression may be linked to adiposity. Therefore, we analyzed TNF-β in an obesogenic LET-induced model of PMOS, this time implanting pellets at 4 weeks of age (peripubertally) (mouse cohort 3) (**Fig. 5A**) (*18*). We confirmed, as previously published, that cohort 3’s peripubertal LET implantation in C57BL/6 mice results in elevated LH and T (**Fig. 5B-C**) after 5 weeks, concomitant with increased weight gain and impaired glucose tolerance (**Supplemental Fig. 8A-B**). We observed a trend towards increased serum TNF-β in LET-treated mice and found that TNF-β positively correlated with LH and fat mass and negatively correlated with lean mass (**Fig. 5D-F, Supplemental Fig. 8C**). To determine whether the observed relationships of adiposity to TNF-β are relevant to women with PMOS, we analyzed serum TNF-β from our previously published age and BMI matched cohort of women with Type A PMOS (human cohort 2) (*38*). Surprisingly, TNF-β was lower in PMOS than controls, even when stratified by BMI. These results are in contrast to previously published data in cohorts that were BMI stratified, but not BMI-matched(*36*) (**Fig. 5G-H**). When we stratified the cohorts by BMI first, we found that TNF-β was increased with overweight/obesity (**Fig. 5H**), indicating that systemic TNF-β is driven by weight, similar in mice. To further understand the role TNF-β might have in PMOS, we performed a moderation analysis (**Fig. 5I**). This analysis determines whether the concentration of TNF-β can influence a correlational relationship between 2 separate variables.

**Figure 5.**
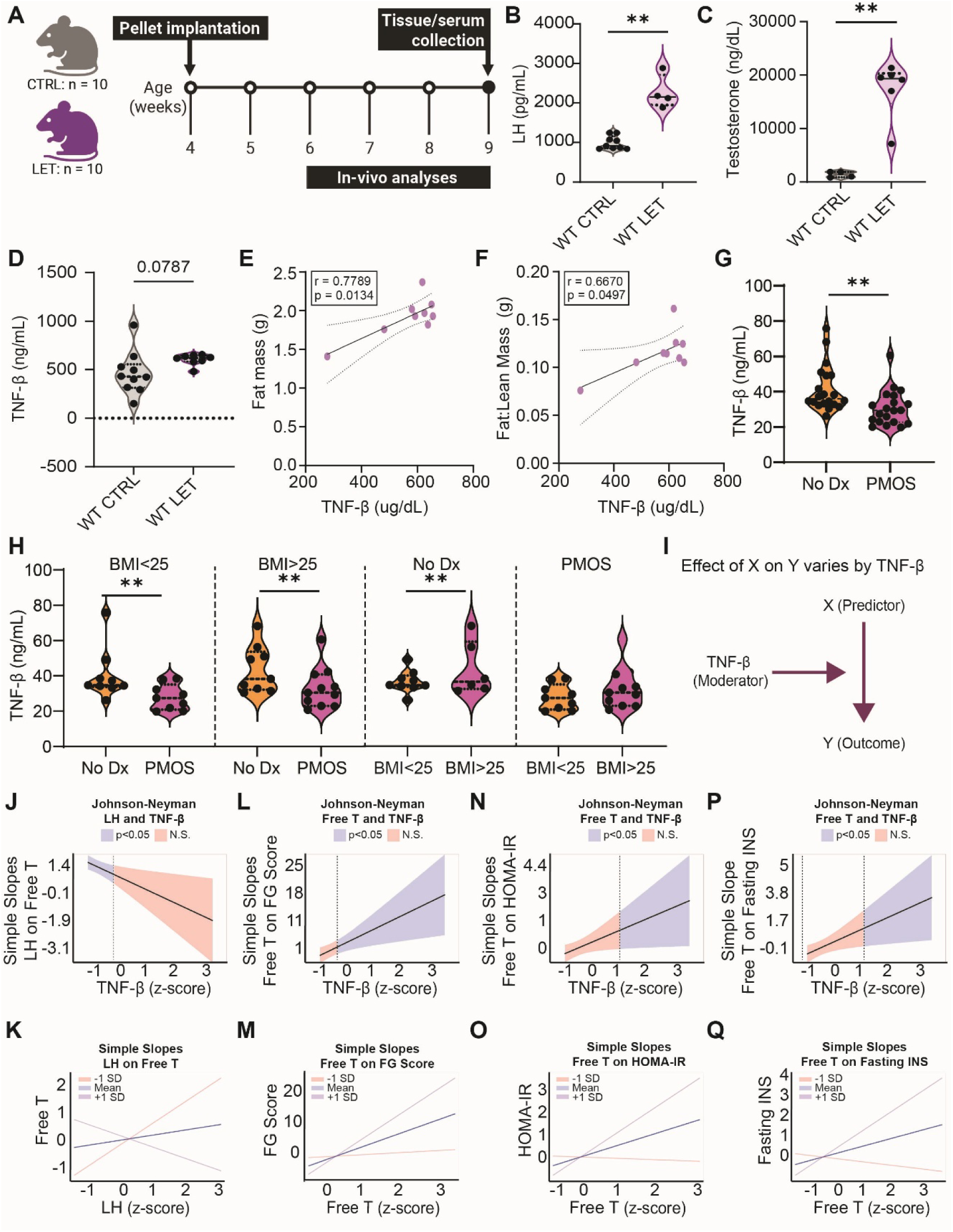
TNF-β is associated with adiposity and modulates the relationship of Androgen to PMOS phenotypes. (**A**) Timeline of cohort 3 letrozole or placebo pellet implantation at week 4 to induce prepubertal model of PMOS-induced symptoms. *In vivo* analyses (glucose tolerance testing, body composition, and estrous cycling) were measured between 6-9 weeks, and sac at 9 weeks included collection serum. (**B-D**) Serum levels of luteinizing hormone (pg/mL) (B), testosterone (ng/dL) (C) and TNF-β (pg/mL) (D). (**E-F**) Correlations of TNF-β and fat mass (E) and fat to lean mass ratio (F) in WT LET cohort. (**G-H**) Levels of TNF-β (ng/mL) in women with and without PMOS (G), stratified by BMI and diagnosis (H). Data are presented as mean ± SEM and analyzed with t-test for parametric data or Mann-Whitney test for nonparametric data. Significance was accepted at p<0.05 and is indicated with an *. (**I**) Moderation analysis structural equation. (**J-Q**) Moderation analysis of TNF-β on the relationship of luteinizing hormone to free testosterone (J and K) free testosterone to Ferriman-Gallwey score (L and M), free testosterone to HOMA-IR (N and O) and free testosterone to fasting insulin (P and Q).

We found that TNF-β significantly moderated the relationship between androgens and hirsutism and also the relationship between androgens and metabolic dysfunction (**Fig. 5 J-Q and Supplemental Figure 8D-E**). Within this cohort, higher TNF-β was associated with stronger androgen-related hirsutism and metabolic dysfunction measures.

### TNF-**β** promotes androgen-independent AR nuclear localization and AR splice variant expression in pituitary/gonadotropic cells

The moderation analyses indicated that TNF-β may modulate androgen responses. To determine whether TNF-β had a direct impact on androgen receptor (AR) activation, we treated dispersed ovary and pituitary with dihydrotestosterone (DHT, a potent androgen) or TNF-β. We analyzed AR translocation by flow cytometry using a specialized cell permeabilization protocol with a cytosolic protein release step (*39*). Cytosolic protein release allows detection of nuclear protein by flow cytometry. We confirmed the validity of this approach by demonstrating that DHT induces AR translocation in the gonadotropic cell line LβT2 (**Supplemental Fig. 9A- C**). We found no effect of TNF-β on AR translocation in ovarian cultures (**Supplemental Fig. 9D-F**). In contrast, TNF-β induced AR translocation in non-immune pituitary cells (**Fig. 6A-C**).

**Figure 6.**
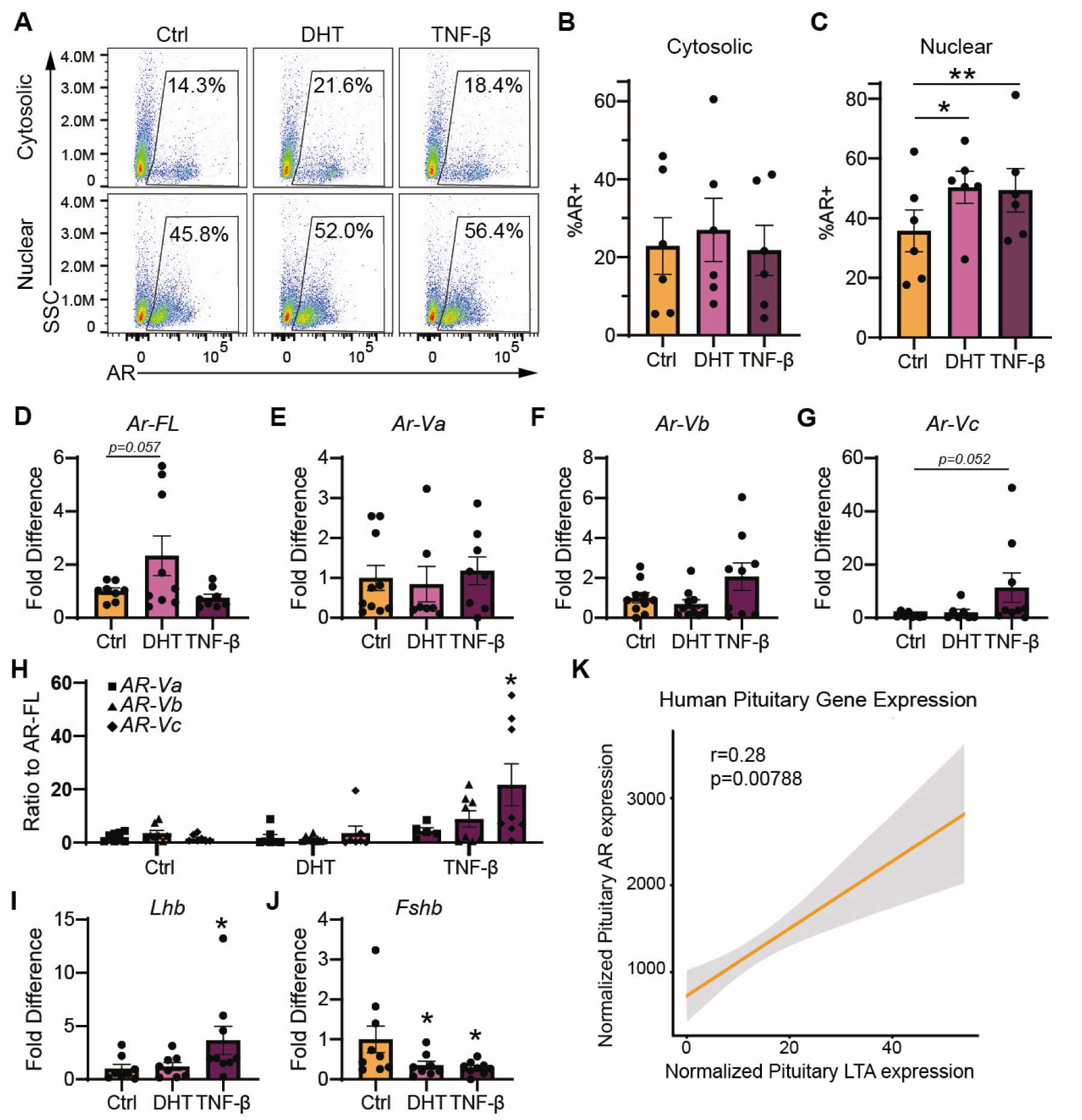
TNF-β induces expression of AR splice variants in LβT2 cells. **(A**) Flow cytometry androgen receptor translocation within nuclear and cytosolic compartments of non-immune pituitary cells treated with DHT and TNF-β. (**B-C**) Quantification of flow cytometry. (**D-G**) Fold increase in mRNA of *Ar-FL, Ar-Va, Ar-Vb*, and *Ar-Vc* in LβT2 cells treated with DHT and TNF-β. (**H**) Ratio of the relative expression of Ar splice variants to full length androgen transcript in LβT2 cells treated with DHT and TNF-β. (**I-J**) Fold increase in mRNA of *Lhb, Fshb,* in LβT2 cells treated with DHT and TNF-β. Data are presented as mean ± SEM and analyzed with one- or two-way ANOVA with a post-hoc Dunnett’s test compared to control. (**K**) Correlation of pituitary LTA expression with pituitary AR expression from the human GTEx database.

TNF-β induction of AR translocation in the absence of androgen indicates an alternative mechanism to AR activation. Androgen-independent AR activation has been extensively characterized in castration-resistant prostate cancer, where AR splice variants lacking the ligand-binding domain exhibit constitutive transcriptional activity in the absence of androgens (*40*).

Given that TNF-β induced androgen-independent AR translocation, we hypothesized that TNF-β promotes expression of AR splice variants in gonadotropic cells. We treated LβT2 cells with DHT or TNF-β and performed qPCR using primers designed to detect full-length AR (AR-FL) and specific AR splice variants endogenous to mice (*41*). TNF-β significantly increased expression of *Ar-Vc* but not transcripts of the full-length receptor or other splice variants (**Fig. 6D-G**). Importantly, we found that the ratio of *Ar-Vc* splice variant to the full-length *Ar* was significantly increased, consistent with TNF-β specifically promoting alternative splicing rather than general upregulation of AR transcription (**Fig. 6H, Supplemental Fig. 9G**). Concurrent with induction of splice variants, TNF-β increases *Lhb* mRNA while reducing *Fshb* mRNA (**Fig. 6I-J**), a direction consistent with the altered gonadotropin profile observed in PMOS-like endocrine phenotypes. These effects are independent of general cell transcriptional activation as DHT and TNF-β equally modulated the immediate early gene *Egr1* (**Supplemental Fig. 9H**). Finally, *LTA* transcript (TNF-β) is correlated with AR in human pituitary (**Fig. 6K**). Collectively, these findings suggest that TNF-β promotes androgen-independent regulation of AR.

## DISCUSSION

In this study, we used a systems genetics immune screen to demonstrate that variation in memory CD4 T cells and TNF-β is associated with LET treatment. By knocking out functional T cells, a major source of TNF-β, we provide evidence for T cells being necessary for the pathological elevation of LH in PMOS-like conditions. The major findings of this study provide a rationale for further investigation of immune contributions to PMOS-associated endocrine dysfunction.

First, we demonstrate that by varying genetic background, we can recapitulate broader presentations of PMOS-like phenotypes. Several BXD strains including BXD124 and BXD60 in addition to the C3H/Hej, DBA, and NOD strains developed phenotypes similar to PMOS type A, the most common amongst the mouse strains (*42*). A/J and BXD125 responded to LET treatment with phenotypes similar to type B PMOS while two BXD48 and BXD79 responded to LET treatment similar to phenotypes of PMOS type C. BXD77 was the only strain that resembled a type D phenotypic presentation of PMOS in response to LET treatment. The ability to generate this diversity of phenotypes in response to LET treatment increases the ability to discover mechanisms common for PMOS phenotypes. There are several animal models of PMOS including the pre-pubertal and adult LET paradigms used in this study. Other models include the administration of dihydrotestosterone (DHT, a non-aromatizable androgen), perinatal and prepubertal exposure to anti-Mullerian Hormone (AMH) among others (reviewed in (*43*)). These models have varied presentation of PMOS-like phenotypes and now by using a genetic diversity panel in the present study, we can capture this same type of diversity using a single paradigm.

The global immune cell composition in the spleen and lymph nodes did not change with LET treatment across the diverse cohort, consistent with published reports showing that circulating immune cell populations are not dramatically altered in PMOS (45). This suggests that changes in immune cell function and cytokine output, rather than immune cell abundance, may be more relevant to PMOS pathophysiology. Thus in this study, we derived cytokine profiles from stimulation of splenic immune cells. Spleen-derived cytokine profiles may not be directly comparable to human serum-derived cytokine profiles because circulating cytokines are indicative of spillover from undetermined tissue sites, while *in vitro* generated cytokine profiles are indicative of the specific tissue (*44*). Blood, though easily accessible, is not the site of immune cell function, therefore mechanistic insight from serum cytokines is limited. This limitation is due to circulating cytokines not being representative of tissue cytokine production, with the exception of strongly inflammatory processes (*45–47*). The apparent discrepancy between elevated *LTA* mRNA in PBMCs (Figure 4I) and lower circulating serum TNF-β protein in women with PMOS (Figure 5G) likely reflects the distinction between cell-intrinsic transcriptional upregulation and systemic protein spillover into circulation. Immune cells in women with PMOS may be primed to express and locally release TNF-β at tissue sites of inflammation, without a proportionate increase in detectable serum protein levels. This compartmentalization is well-documented in other inflammatory conditions and supports the need for tissue-level immune profiling in future studies. Several diseases show discrepancy in serum cytokines vs local tissue cytokine induction and in PMOS. For example, serum cytokine signatures are vastly different from follicular fluid cytokine profiles (*48*). Given that gene expression of *LTA* correlated with changes in gene signatures in the adipose and that serum TNF-β increases with increased adiposity, it is not surprising that in our study, serum TNF-β in a carefully BMI matched cohort was not increased in PMOS, contrary to published data demonstrating increased circulating TNF-β only in lean women with PMOS who were not BMI matched (*36*). To support targeting T cell derived TNF-β in PMOS, it will be important to determine the tissue and cell source of TNF-β in PMOS through comprehensive immune profiling from PBMCs and tissue samples from women with and without PMOS across all subtypes in future studies. Efforts in this area are ongoing (*48*).

There is strong precedent that targeting immune dysfunction is beneficial for PMOS. For example, women with PMOS who take metformin, known to have immunomodulatory properties in addition to reducing hepatic gluconeogenesis, experience improvement of ovulatory function (*49–52*). Because metformin has pleiotropic metabolic and immunomodulatory effects, its clinical benefit is consistent with the possibility that immune pathways contribute to reproductive phenotypes in PMOS. Given there has been no explicit target for immune intervention in PMOS, clinical trials directly testing immune therapy for improvement of PMOS symptoms has not been justified. Our study provides a rationale for further testing whether specific T cell subsets or T cell-derived cytokines contribute to PMOS-associated endocrine dysfunction through induction of androgen independent splice variants. We were able to detect increased transcripts of TNF-β in PBMCs from women with PMOS. While the TNF-β mRNA elevation in PBMCs in women with PMOS is intriguing, these data should be interpreted cautiously because the human PBMC cohort in this study is small, particularly for the controls (n=5). These human data set the stage for more definitive analyses in larger cohorts.

While all direct experimental evidence demonstrates that testosterone (T) suppresses LH secretion (*53–59*), women with PMOS exhibit both elevated T and increased LH. It is possible that in PMOS, testosterone feedback to reduce neuroendocrine LH secretion is nonfunctional or disrupted, permitting abnormally high LH in the face of high androgens. The most striking finding in our study was that the abnormally increased LH in the LET mouse model was not observed in TCRα KO mice, despite their increased testosterone due to aromatase inhibition. Our data suggest that T cells may be involved in the disruption of this feedback mechanism, since LH was no longer elevated in LET TCRα KO females (lacking T cells), despite elevated testosterone. Together with data demonstrating that TNF-β positively regulates AR splice variants and LH, these results raise the possibility that T cell-derived cytokines could contribute to altered sex steroid feedback in PMOS-like conditions. Thus, it is possible that a lack of T cells permitted testosterone negative feedback inhibition on LH to occur. This intriguing possibility requires future detailed investigation to uncover mechanisms of T cell action and how this intersects with testosterone feedback signaling on neuroendocrine targets. For example, adoptive transfer of memory T cells from LET female mice to control mice could test whether T cells impede testosterone negative feedback. Future studies can also employ the peripubertal LET paradigm in TCRα KO mice to see if those females are also protected from developing metabolic dysfunction, in addition to reproductive protection (*18*). A key limitation of the estrous cycle assessment in this study is that monitoring was limited to one cycle (4-8 days) during the final week of LET treatment, which may not capture the full spectrum of cycle disruption. Future studies directed at impacts of immune function on cyclicity should assess cycling for a minimum of 12 consecutive days to comprehensively characterize cycle arrest patterns.

This study has several limitations, some of which are noted above in the relevant experimental sections. The causal evidence for T cells derives from the LET-induced PMOS-like mouse model and requires validation in additional models and human tissues. TCRα KO mice support a role for functional αβ T cells in LET-induced LH elevation, but this approach does not identify the responsible T cell subset or exclude developmental or compensatory effects of lifelong TCRα deficiency. TNF-β should also be considered a candidate mediator, as direct in vivo blockade or receptor-specific testing was not performed. Human analyses are limited by small cohort size, bulk PBMC measurements, and cross-sectional serum data, which prevent identification of the cellular source of TNF-β or causal inference. Additional in vivo studies are needed to determine whether TNF-β acts directly on gonadotropes to regulate LH secretion in PMOS-like conditions.

Despite these limitations, we identify TNF-β as a candidate target meriting further investigation in the context of PMOS-associated immune dysfunction. Before this work can be translated into a feasible treatment for women with PMOS, there are several questions that remain to be answered. First, which specific T cells are mediating the PMOS-like phenotypes must be determined. Our present cytokine profiling implicates a potential Th17 mechanism, consistent with Th17 cell secretion of TNF-β (*60*). Finally, rodent models of PMOS are limited, given there is no natural diagnoses of PMOS in rodents (*61*). To truly translate this work to the clinic, mechanisms of T cell disruption of androgen feedback would need to be performed in primary human pituitary followed by intervention studies performed across strains representing all four subtypes of PMOS with robust sample size.

## MATERIALS AND METHODS

### Animals

All animal procedures were carried out in accordance with relevant guidelines and regulations. Experiments performed at the University of California, Irvine were approved by the University of California, Irvine Institutional Animal Care and Use Committee under protocols AUP-22-102 and AUP-24-068. Animal procedures performed at the University of California, San Diego were approved by the University of California, San Diego Institutional Animal Care and Use Committee under protocol S14011. Mice were maintained under a 12-hour light/dark cycle with ad libitum access to food.

#### Mouse Cohort 1. Systems genetics screen

Female inbred strains were acquired from Jackson Laboratories, and BXD strains were provided by Robert W. Williams and David G. Ashbrook (**Supplemental Table 1**). Our PMOS model was established using a protocol adapted from prior studies (*18, 29*). Female mice, aged 9-10 weeks, received subcutaneous implants of letrozole (PMOS) or placebo pellets (3mm diameter; Innovative Research of America) at the start and after 3 weeks, ensuring a continuous release of 50 μg/day of letrozole over 6 weeks. Letrozole-treated and control mice were housed separately, with no more than four mice per cage.

#### Mouse Cohort 2. TCR alpha KO mice

TCR alpha KO mice were acquired from Jackson Laboratories. Placebo and LET pellets were inserted at 9 weeks of age as illustrated in Figure 1. Two wildtype sentinel mice were included with the cohort to ensure the current batch of LET pellets were inducing PMOS-like symptoms. The mice underwent routine body composition analysis, echocardiography, glucose tolerance tests, estrous cycle assessments, and weekly weight measurements.

#### Mouse Cohort 3. Obesogenic PMOS mouse model

Wildtype C57BL/6 female mice were acquired from Jackson Laboratories. Placebo and LET pellets were inserted at 4 weeks of age (peripubertally) as illustrated in Figure 5. For all cohorts, mice were euthanized using 2.5% isoflurane delivered via a precision vaporizer, followed by cervical dislocation. Blood was collected via cardiac puncture immediately following euthanasia. Metabolic and reproductive tissues were collected, immediately frozen in dry ice, and stored at −80°C. One ovary per mouse was fixed in 4% paraformaldehyde at 4°C overnight, preserved in 70% ethanol, and processed for histology. Serum was collected after coagulation at room temperature for 1 h (2,000 x g for 10 min at 4°C) and stored at -80°C for subsequent analyses.

### Ethics approval and consent for human samples

All methods involving human participants and human-derived samples were carried out in accordance with relevant guidelines and regulations. For Human Cohort 1, peripheral blood samples from women with and without PMOS were collected at Poznań University of Medical Sciences under a protocol reviewed and approved by the Bioethics Committee at Poznań University of Medical Sciences, Poland (approval no. KB-1227/17). Written informed consent was obtained from all participants prior to sample collection. For Human Cohort 2, de-identified plasma samples and associated clinical metadata were obtained from the Androgen Excess Biorepository as part of a previously published study (*38*). That study was approved by the institutional review boards at both the University of Alabama at Birmingham and Cedars-Sinai Medical Center, and written informed consent was obtained from each participant.

### Human primary samples

#### Human Cohort 1

Human peripheral blood was collected from Dr. Beata Banaszewska’s patient list of women with PMOS (n=15) or without PMOS (n=5). Dr. Beata Banaszewska carried out all clinical diagnostics and measurements of her patients. Peripheral blood mononuclear cells were isolated using a density gradient media and cryopreserved and shipped overnight in dry ice from Poznan University of Medical Sciences, Poland to the University of California, Irvine where they were stored in liquid nitrogen for downstream application.

#### Human Cohort 2

Human plasma samples and associated clinical metadata were obtained from the Androgen Excess Biorepository as part of a previously published prospective study of women evaluated for androgen excess at the University of Alabama at Birmingham and Cedars-Sinai Medical Center between 1987 and 2010 (*62*). Fasting follicular-phase blood samples were collected, processed, and stored in the biorepository for downstream analyses. Clinical evaluation, diagnostic classification, and sample collection were performed as previously described (*38*). PMOS diagnosis was defined using the 1990 NIH criteria. For the present analysis, a cross-sectional case-control subset consisting of 40 women with PMOS and 40 matched controls was selected. Participants were matched for race, age, and BMI and included equal representation of White and Black women.

### Body weight, body composition, and glucose tolerance test

Body weights were measured at the beginning of each week, and body composition measurements were taken using the EchoMRI™ Whole Body Composition Analyzer on the 5th week of the protocol. Intraperitoneal (ip) glucose tolerance tests (GTT) were performed on the 5th week in a conscious state. Briefly, mice fasted for 6 h were ip injected with 1 g of glucose/kg body mass, and glucose levels were determined from the tail-tip using a hand-held glucometer (Accu-Check Guide) in the basal state and 15, 30-, 45-, 60-, and 90-min following glucose administration.

### Estrous Cycle Assessment

Due to the number of *in vivo* assessments carried out on the mice, and to reduce stress on the animals, only one estrous cycle (4-8 days, standard for the LET paradigm) was assessed during the final week of the protocol (6th week), which allowed us to determine if the last cycle was normal (normal progression diestrus-proestrus-estrus-meta estrus) or not (arrested cycle). Estrous cycle assessment during the final week of LET treatment is standard for this paradigm, given the consistent and reproducible arrest in diestrus on the BL6 J background with treatment. The estrous cycle stage was determined by light microscopic analysis of smears from the vaginal lavage by 2 independent, blinded experts. Classifications were compared and discrepancies were resolved through consensus. Proestrus was defined by the presence of mostly nucleated and some cornified endothelial cells, estrus as mostly cornified cells, metestrus as some cornified endothelial cells and mostly leukocytes, and diestrus as primarily leukocytes. If uncertainty remained, the machine learning model ODES was used as a supportive tool to assist in the final classification (*63*).

### Ovarian morphology

To roughly estimate ovarian follicular populations, partial follicular counting and classification were performed (approximately ¼ per ovary per mouse) in ovarian serial sections (5 um/section). In brief, fixed ovarian tissue was consecutively cut, placed on gelatin-coated slides (Biobond, British Biocell International, Cardiff, UK), air dried for 2 h, and fixed for 5 min in acetone at 4 C. Subsequently, consecutive sections from each ovary were washed in PBS (137 mmol/l NaCl, 2.7 mmol/l KCl, 4.3 mmol/l Na2 HPO4.7H2O, 1.4 mmol/l KH2PO4, and pH 7.3) and stained with hematoxylin and eosin (DAKO Corporation, Carpinteria, CA, USA) for histological analysis. Histological serial sections were independently analyzed by three of the authors, and ovarian follicles were classified and quantified. The number of follicles was counted at a regular interval, and the total number of follicles per ovary was estimated using a multiplication factor based on follicle class and sampling fraction (*64*). Cystic follicles were fluid-filled structures outlined by a single layer of granulosa cells and hypertrophic theca cell layer. Primordial, primary, secondary, and antral follicles were counted and classified as described previously (*65*). Then, follicles were grouped into primordial, small follicles (grouping primary, secondary, and small antral follicles), and large follicles (Graafian/preovulatory follicles). Follicular atresia was also quantified, and atretic follicles were defined as follicles with >5% of the granulosa cells having pyknotic nuclei.

### Serum terminal assays of hormonal levels

Serum hormonal levels of luteinizing hormone (LH) (UVA in house protocol, sensitivity 0.016 ng/mL, range 0.016-4 ng/mL), follicle-stimulating hormone (FSH) (UVA in house protocol, sensitivity 0.016 ng/mL, range 0.016-8 ng/mL), estradiol (E) (Alpco, 55-ESTRT-E01, sensitivity 2.5pg/mL, range 5-1280 pg/mL), and testosterone (T) (ILB, IB79174, sensitivity 0.006 ng/mL, range 0.1-25 ng/mL) (**Figure 1**) were assessed by The University of Virginia (UVA) Center for Research in Reproduction Ligand Assay and Analysis Core. Insulin levels in serum were measured with the Mouse Insulin ELISA (Alpco, catalog 80-INSMS-E01). Due to reduced sample availability, serum testosterone (**Figure 3**) was measured via the Multi-Species Hormone kit (MSHMAG-21K) and testosterone flexing pack (SPRCA1825). The testosterone capture antibody exhibits 100% reactivity to Testosterone and 16% cross-reactivity to 5-alpha-DHT. Briefly, 50 µL of serum was vortexed with 75 µL acetonitrile and incubated for 10 minutes at room temperature. The sample was then vortexed again for 5 seconds, then centrifuged at 17,000 x g for 5 minutes. 100 µL of supernatant was transferred into new Eppendorf tubes. The samples were dried by Speed Vac, followed by reconstitution with 40 µL Luminex Assay Buffer before being analyzed by multiplex assay. Finally, human LH was measured by Luminex assay (HPTP1MAG-66K, Millipore, MILLIPLEX® Human Pituitary Panel 1, sensitivity 1.12 pg/mL, std curve range 122-500,000 pg/mL).

### Data Scaling and Normalization

To address the systematic differences between the Luminex assays and ELISAs used to measure Testosterone, LH, and FSH in Figure 3, we employed a linear scaling method based on placebo and LET treated WT C57BL/6 mice female mice. C57BL/6 mice on the LET experimental paradigm have robust, consistent, and repeatable hormone outcomes (*18, 19, 29*). The scaling was performed to adjust the measurements from the Luminex (Assay 2) to be comparable with those from the University of Virginia (UVA) Center for Research in Reproduction Ligand Assay and Analysis Core (Assay 1). For each analyte, we calculated the mean (μ) and standard deviation (σ) of the measurements from both assays. We then applied the following formula to scale the Assay 2 data:

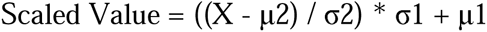

Where X is the original measurement from Assay 2, μ2 and σ2 are the mean and standard deviation of the Assay 2 C57BL/6 dataset, and μ1 and σ1 are the mean and standard deviation of the Assay 1 C57BL/6 dataset. This transformation adjusts the Assay 2 WT Placebo data to have the same mean and standard deviation as the Assay 1 WT Placebo data, allowing for direct comparison between the two datasets. The scaling was performed using Microsoft Excel. Both the original and scaled data were retained for reference.

### Variance partitioning analyses (heritability)

Mixed effect models were used to estimate genetic and PMOS contributions to traits, as shown previously (*66*). In brief, variances were partitioned using a linear random slope model (nlme R Package) with treatment and strain as random effects. The proportion of the total variance explained by strain was scaled to explain a portion of the residual variance from the previous model as a strain-by-diet interaction. Linear modeling was performed using lme functions from the R package lme4 version 1.1-25. Variance plots were generated using the R package ggplot2 version 3.3.2.

### Splenocyte stimulation and cytokine multiplex immunoassay

At the end of the 6-week treatment, the spleen of the mice was collected. The spleen cells were mechanically dissociated and stimulated (200,000 cells/200 µL) with either mouse T-activator CD3/CD28 dynabeads (1 bead per cell; ThermoFisher) or lipopolysaccharide O111:B4 (conc.; Ebioscience) for 40 hours at 37°C. The conditioned media from the stimulated splenocytes for each mouse were collected and stored at -20°C. A total of 40 cytokines and chemokines were quantified using the Milliplex MAP mouse Th17 magnetic bead panel and the Milliplex MAP mouse cytokine/chemokine magnetic bead panel (Millipore Sigma) according to the protocol from each respective kit. The analyte concentrations were measured using an Intelliflex instrument (Luminex) and the Belysa software (1.2). Two separate datasets of cytokine quantifications were generated based on whether the supernatant originated from splenocytes activated with either CD3/28 dynabeads or LPS.

*L*β*T2 cell culture*

The female C57BL/6 mouse-derived LβT2 gonadotrope cell line was maintained in high-glucose (4.5 g/l) HEPES-buffered DMEM supplemented with penicillin/streptomycin and 10% fetal bovine serum (FBS: FB-11, Omega Scientific, CA) at 37 °C in a humidified atmosphere of 5% CO_2_ (*67*). To test the effects of DHT and TNF-β, LβT2 cells were seeded at 2 × 10^5^ cells per cm^2^, cultured for 24 h, and pretreated with serum-free and phenol red-free DMEM for 12–16 h prior to treatment. Cells were treated with dihydrotestosterone, 10 nM DHT (Sigma-Aldrich A8380-1G,) or 10 ng/mL TNF-β (R&D Systems 211-TBB-010/CF,) followed by Flow cytometry or qPCR.

### Staining and Flow Cytometry

All mice inguinal lymph nodes were enzymatically dissociated using 0.25% collagenase for 30 minutes, inhibited with 0.5M EDTA and then mechanically dissociated using a 50μM strainer washed with FACS buffer. While enzymatic digestion can affect certain surface markers, this protocol was selected to achieve adequate single-cell suspensions from lymph node tissue while processing high numbers of tissues. Mice spleens were mechanically dissociated using a 50uM strainer and incubated with red blood cell lysis for 5 minutes before being washed with FACS buffer. Cells were aliquoted into 96-well plates for viability staining and the staining of the 18 markers (**Supplemental Table 2**). Flow cytometry was performed on Cytek Northern Lights Spectral Cytometer. Gating was performed using Fluorescence Minus One (FMO) controls for each fluorophore to determine positive/negative boundaries. All gates were set using the FMOs and applied consistently across all samples. To correct for batch effects across several flow cytometry runs, data was processed in OMIQ using CytoNorm and presented as normalized counts. Normalized counts represent the number of cells in each identified immune cluster normalized to total cell input to account for differences in cell recovery across samples.

### AR translocation Flow Cytometry Assay

Mouse pituitaries and ovaries were mechanically and enzymatically dissociated in collagenase IV (Gibco) for 20 min at 37°C in phenol red-free DMEM (Gibco). Following dissociation, cells were plated and allowed to adhere overnight, followed by 12-hour serum starvation in phenol red-free media before treatment with 10 nM DHT or 10 ng/mL TNF-β for 1 hour. Following treatment, cells were collected and AR translocation was analyzed by flow cytometry using a differential permeabilization protocol from Brittain and Gulnik (*39*). Cells were stained with Zombie NIR viability dye (Biolegend, 1:3200) according to manufacturer’s instructions. After Fc block (Fcx TrueStain, Biolegend) and fixation, cells were stained for CD45 and stained with Androgen Receptor (**Supplemental Table 3**) Polyclonal Antibody conjugated to FITC (ANDR-FITC, Thermo Fisher Scientific, 1:5000) and Mouse CD45 Alexa Fluor® 405-conjugated Antibody (Clone 30-F11, Catalog # FAB114V, R&D Systems, 1:500) using the overnight protocol (*68*). The %AR positive cells was reported of live, single, CD45- cells.

### Moderation Analysis

Moderation analysis was performed using JMP Pro statistical software (SAS Institute Inc., Cary, NC) to determine whether TNF-β concentration influences the correlational relationship between androgens and PMOS clinical outcomes. Prior to analysis, all continuous variables were standardized using z-score transformation (mean-centered and scaled by standard deviation) to ensure comparability across different measurement scales. First, Pearson correlation analyses were conducted to identify significant bivariate relationships between clinical and biochemical variables in the cohort of women with PMOS (n=20 for androgen, n=10 LH) and without PMOS (n=20 for androgen, n=14 LH). Variables demonstrating significant correlations (p<0.05) were selected for subsequent moderation analysis. Moderation analyses were then performed with testosterone or LH as the independent variable (predictor), hirsutism score or metabolic dysfunction as the dependent variable (outcome), and serum TNF-β concentration as the moderator. The moderation model tested the interaction effect between testosterone and TNF-β on each outcome variable. The model was specified as:

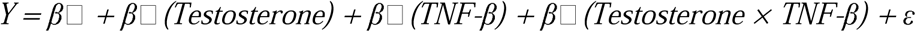

where Y represents the outcome variable (hirsutism or metabolic dysfunction), β□represents the interaction term (moderation effect), and ε represents the error term. A statistically significant interaction term (p<0.05) indicated that TNF-β significantly moderated the relationship between testosterone and the outcome variable.

### RNA isolation, reverse transcription and quantitative polymerase chain reaction

Cryopreserved peripheral blood mononuclear cells from women with PMOS (n = 15) or without PMOS (n = 5) were thawed in a 37°C water bath and washed with 10mL of RPMI 1640 media supplemented with 10% FBS. Cells underwent RNA isolation via Invitrogen’s TRIzol Reagent protocol (CAT #15596018). mRNA from DHT or TNF-β treated LβT2 cells was isolated using RNeasy Plus Micro Kit (Qiagen). Complementary DNA was made by reverse transcription of 2 μg total RNA using Applied Biosystems’s High-Capacity cDNA Reverse Transcription Kit and protocol (CAT #4368814). Complementary DNA products were detected using iQ SYBR Green Supermix (Bio-Rad Laboratories) on a CFX Opus 384 Real-Time PCR System. Data were analyzed by the 2ΔΔCt method by normalizing genes of interest (shown below) to *Gapdh*. Primers for genes of interest were designed using NCBI’s BLAST and Primer Designing Tool. Primers for *Ar* splice variants were derived from literature (*41*).

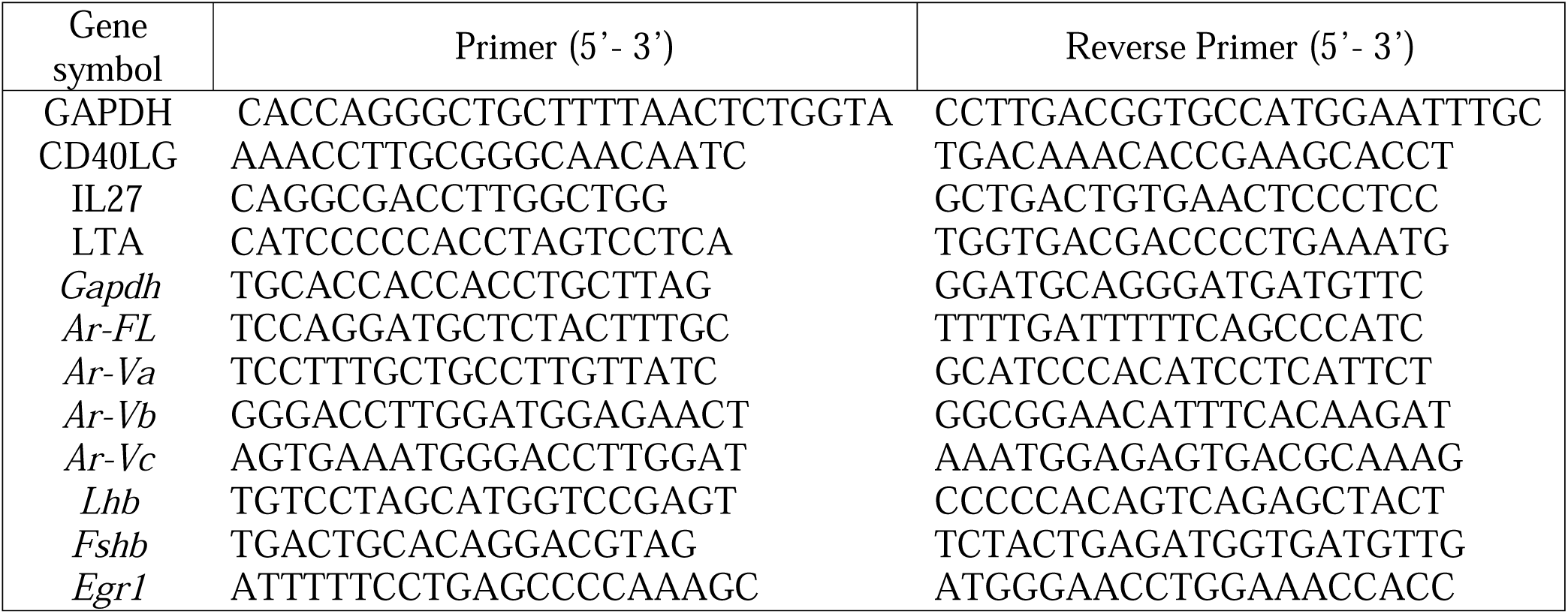

### Partial least squares modeling and principal component analysis

Partial least squares discriminate analysis (PLS-DA) and principal component analysis (PCA) was done for the two datasets of cytokine and flow cytometry quantifications separately using Solo (Eigenvector Research, Inc.). Measurements were assigned as the independent variable, while the discrete regression variable was the treatment classification (LET or control). Two primary latent variables, LV1 and LV2, were used in leave-one-out cross-validation (n=<20) or venetian blinds (n>20). The model was deemed significant in ability to classify with an error rate greater than 70%.

### Correlation analyses

To analyze cytokine-phenotype and genetic correlations corresponding to cross tissue pathways enriched for splenic LTA, we applied established analyses (*69–71*). Specifically, expression of LTA in 31 premenopausal women (age <50) from GTEX v8 (*37*) was correlated with all other genes in indicated tissues using midweight bicorrelation (*72*) and corresponding students regression p-value. Pathway enrichments from these cross tissue correlations were performed using gseGO and enrichR where the bicor coefficents were used as gene weights and mapped to all expressed genes from the indicated tissue. Intrapituitary correlation of LTA with AR was performed in the GTEx online platform gdcat.org. Significance was reported on Benjamini-Hochberg (BH) FDR adjusted p-values.

### Statistical analysis

All the statistical tests were performed in R (4.2.3) and PRISM GraphPad (9.5.1). All continuous data were tested for normality using the Shapiro-Wilk test and visually assessed using Q-Q plots. For normally distributed data Student t-tests were performed in comparisons between the LET and control mice groups. For data that did not meet normality assumptions, Mann-Whitney U test (non-parametric) was performed. To compare population percentages, z-tests were performed. Experimental design with more than 2 groups were analyzed with one-way ANOVA for normally distributed data, followed by a post hoc Dunnett’s test to compare experimental to control. Kruskal-Wallis test was used for non-normally distributed multi-group comparisons. All statistical analysis results with a p-value<0.05 were determined as significant.

## List of Supplementary Materials

Figs. S1 to S9

Tables S1 to S3

## Acknowledgments

We gratefully acknowledge Dr. Robert W. Williams and Dr. David E. James from the University of Tennessee for generously providing the mice used in this study. Their contribution was instrumental to the success of this research. Dr. Nicholas would like to express her sincere gratitude to her postdoctoral mentor Dr. Mark Lawson from the University of California San Diego for his unwavering support in development of the ideas presented in this work, particularly during the preparation of her K99/R00 application. His guidance, mentorship, and encouragement were invaluable in completing this first fully independent project in her laboratory.

## Funding

NIH R00 HD098330 (DN)

NIH R00 HD098330 S1 (NU, DN)

NIH R01 HD095412 (VGT)

NIH T32 HD007203 (JC)

NIH K12 GM068524 (JC)

NIH K99 HD107217 (JC)

NIH DP1 DK130640 (MS)

## Author contributions

Conceptualization: NU, LV, MS, DN

Methodology: NU, LV, GDR, CN, KW, ZDM, SG, JC, JA, BB, RAR, ASK, VGT, BB, EW, AD, MS, DN

Investigation: NU, LV, GDR, CN, KW, ZDM, SG, JC, JK, BB, AD, NN, JA, ASK, VGT, BB, EW, RA, MP, JC, DN

Visualization: NU, LV, GDR, CN, KW, JK, NN

Supervision: NU, LV, CN, KW, RAR, VGT, BB, EW, AD, MS, DN

Writing—original draft: NU, DN

Writing—review & editing: NU, LV, GDR, CN, KW, ZDM, SG, JC, JK, BB, AD, NN, JA, RAR, ASK, VGT, BB, EW, AD, MS, RA, MP, JC, DN

## Competing interests

Authors declare that they have no competing interests.

## Data Availability

All data generated or analyzed during this study are included in this published article and its Supplementary Information files. Additional raw data supporting the findings of this study are available from the corresponding authors upon reasonable request. Publicly available GTEx data analyzed in this study are available through the GTEx Portal: https://gtexportal.org/home/ and https://gtexportal.org/home/downloads/adult-gtex/bulk_tissue_expression. The GTEx Analysis Release V8 dataset is available under dbGaP accession number phs000424.v8.p2 at https://dbgap.ncbi.nlm.nih.gov/home/.

## Supplementary Figures

**Supplemental Figure 1.**
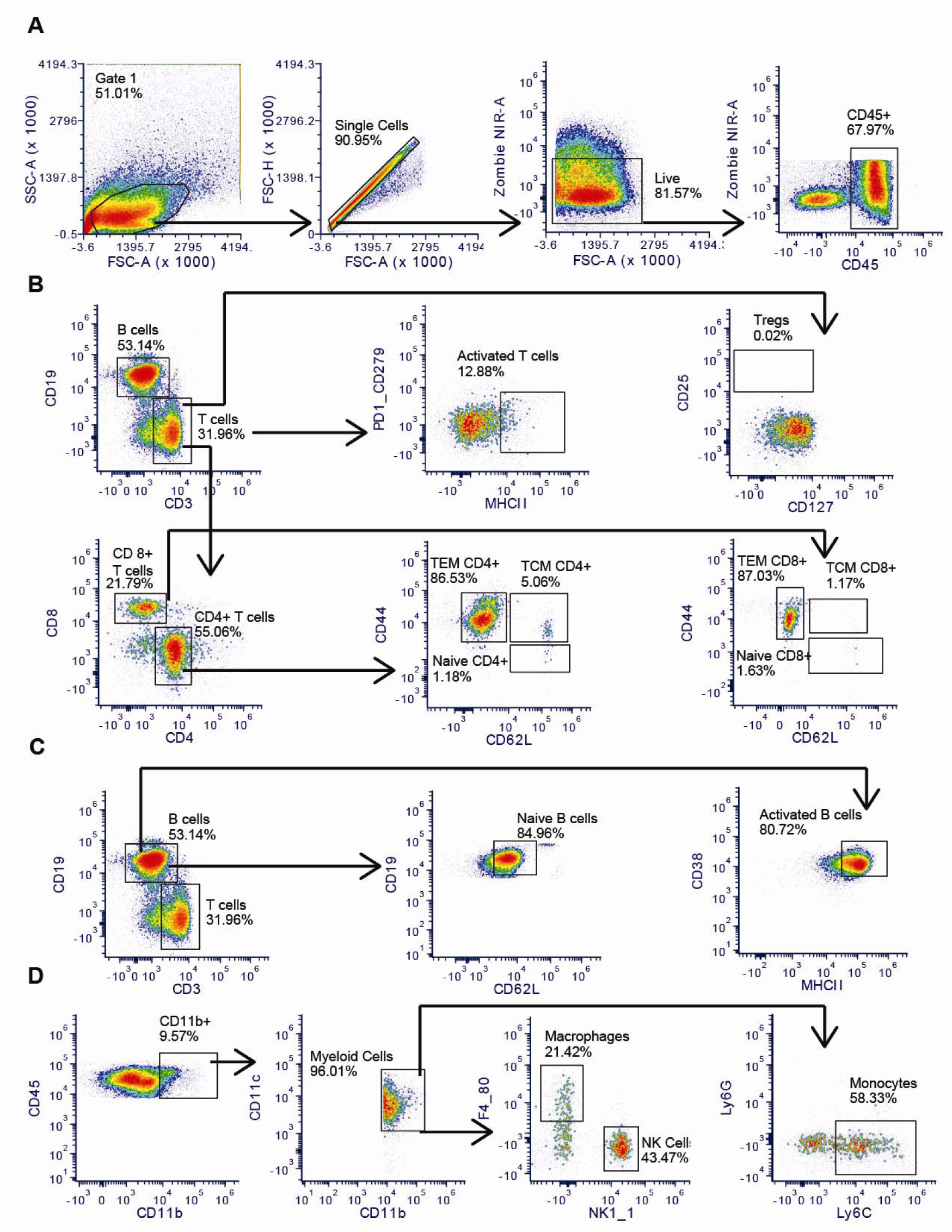
Flow cytometry gating strategy of 18 color antibody immunophenotyping panel. (**A**) Live, CD45+ immune cells, (**B**) activated T cells, T regulatory cells, CD4+ and CD8+ naïve and memory T cells, (**C**) naïve and activated B cells, (**D**) myeloid cells. NK and NKT cells were identified within the CD11b+ myeloid gate using NK1.1 positivity. This gating approach does not capture all NK cells and does not discriminate between NK and NKT cells. Representative LET treated C57BL/6J mouse is shown. FMOs were used for all gates.

**Supplemental Figure 2.**
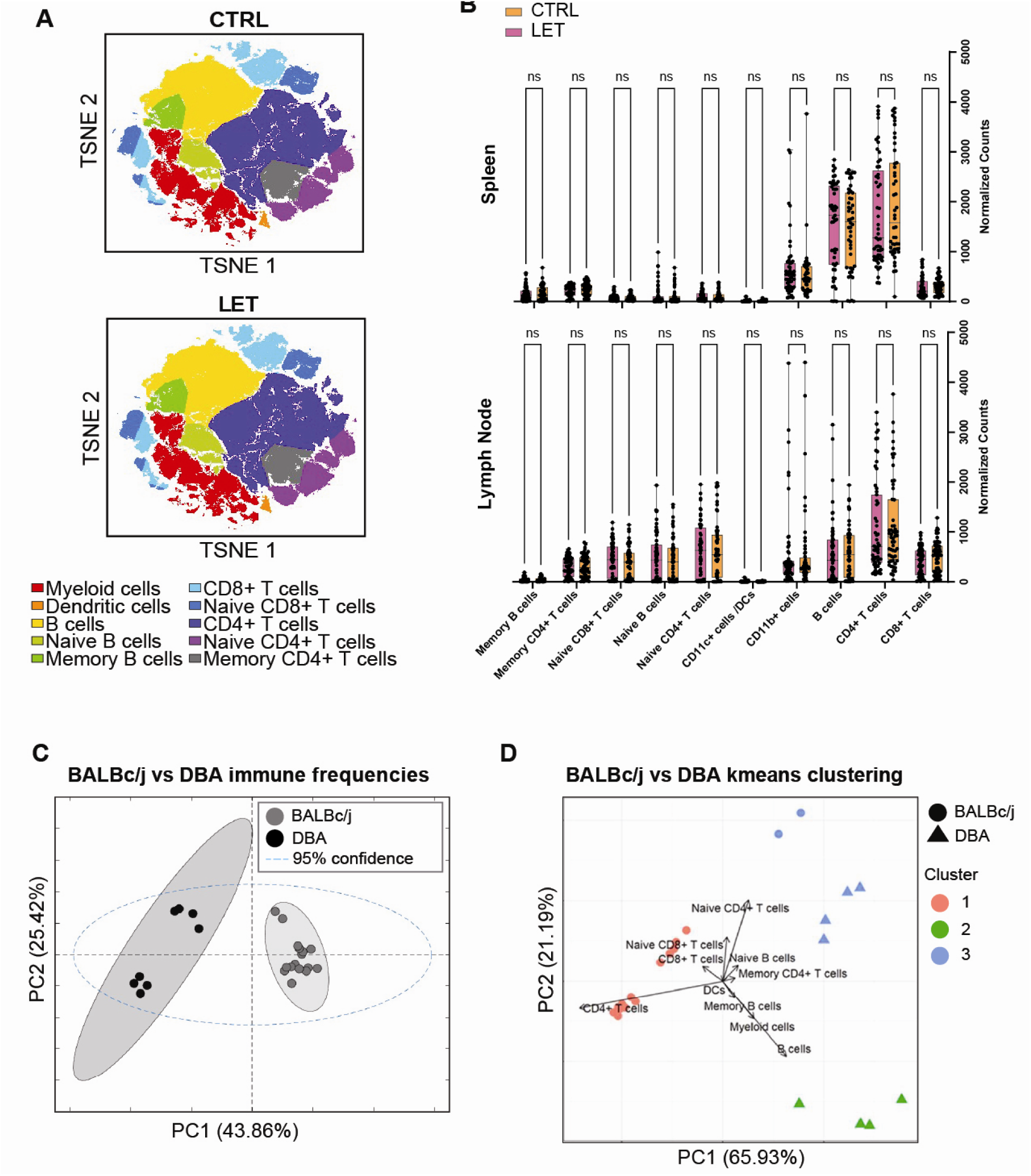
Immune cell frequencies do not shift with LET-treatment. (**A**) TSNE plot of spleen and lymph node immune cell phenotyping by flow cytometry from the genetically diverse CTRL and LET cohort 1. (**B**) Normalized counts (to total cell input) of immune cell clusters in spleen and lymph nodes. Data are presented as mean ± SEM and analyzed with t-test for parametric data or Mann-Whitney test for nonparametric data. Significance was accepted at p<0.05 and is indicated with an *. (**C**) PCA plot of immune frequencies via flow cytometry from BALBc/j and DBA strains. (**D**) PCA plot with K means clustering projection of immune frequencies via flow cytometry from BALBc/j and DBA strains.

**Supplemental Figure 3.**
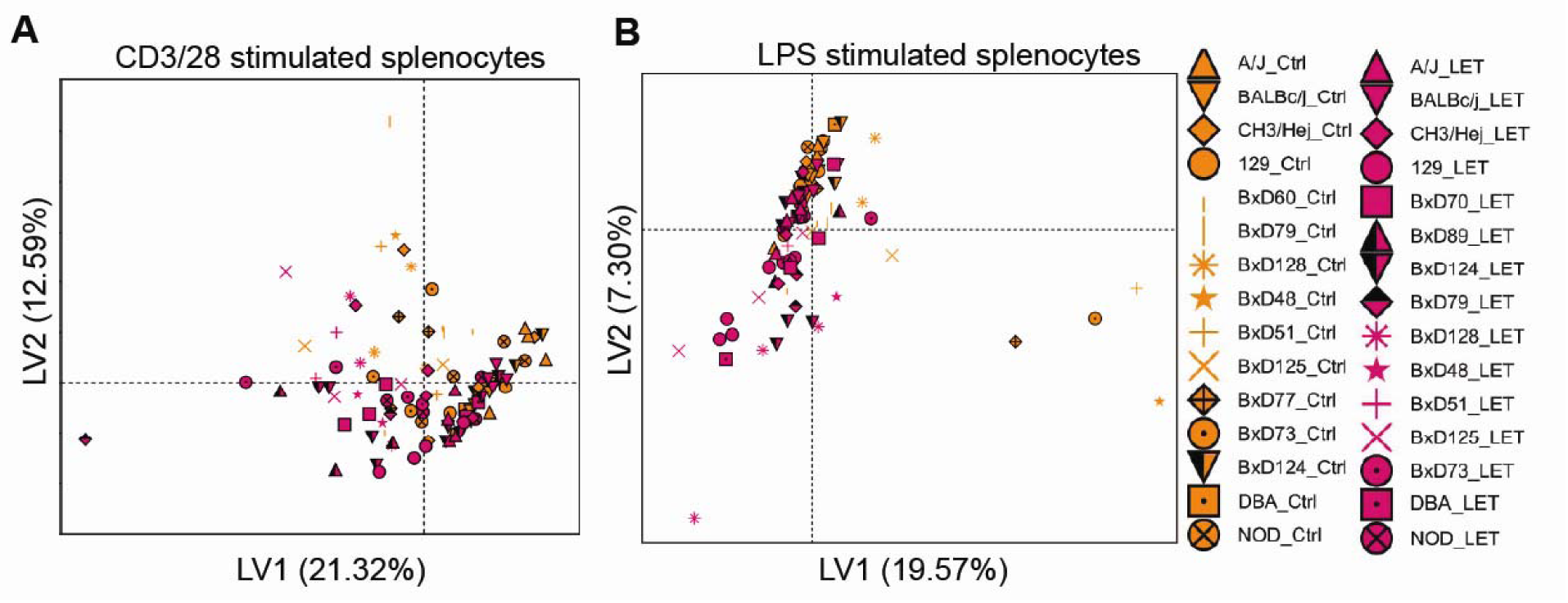
Cytokine profiles vary by strain and are modulated by LET treatment. PLSDA of 40 cytokines measured from the conditioned media of CTRL and LET splenocytes stimulated with anti-CD3/28 (**A**) or LPS (**B**) from cohort 1.

**Supplemental Figure 4.**
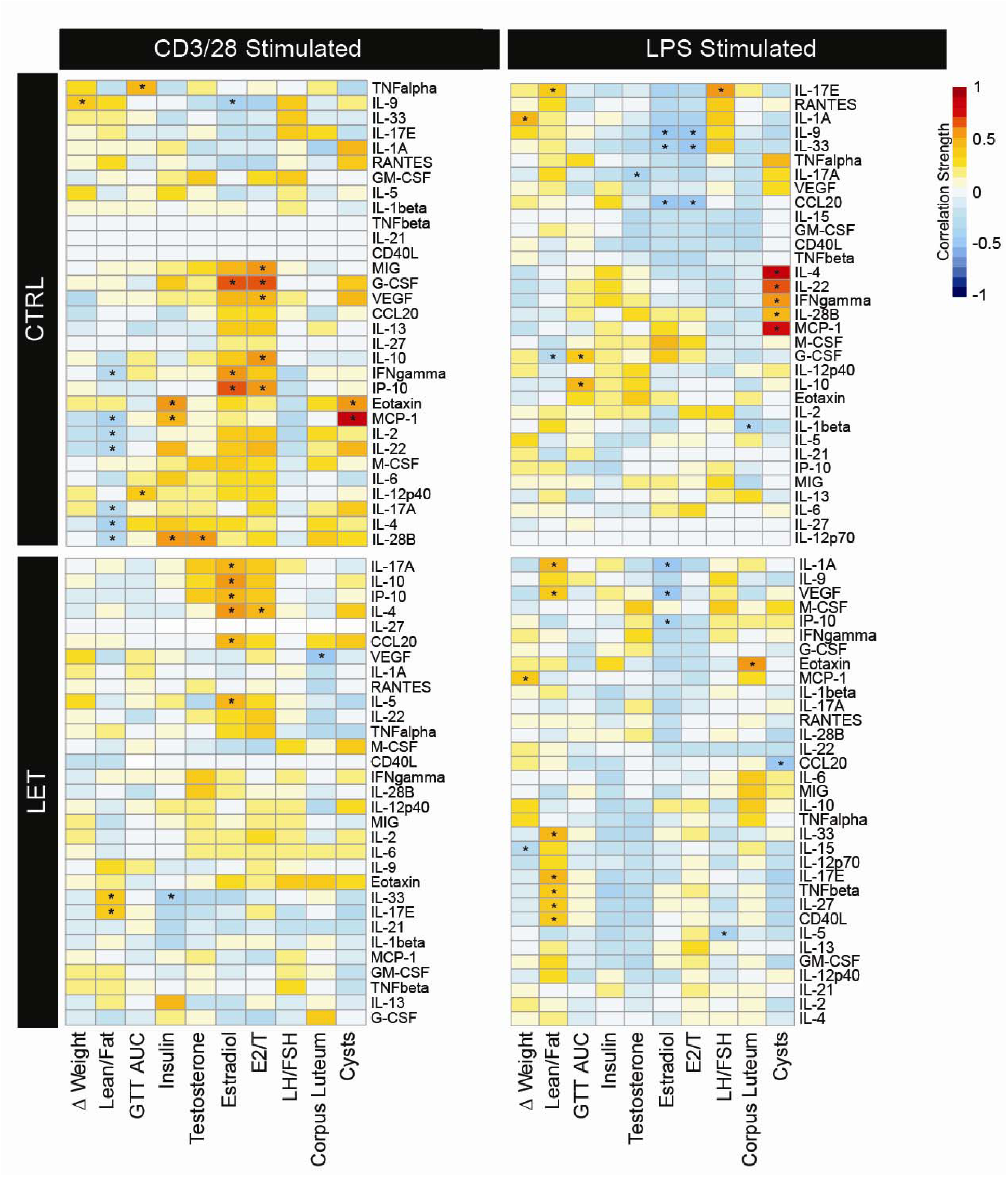
T cell derived cytokines correlate with LET induced PMOS like symptoms. Heatmap depicting correlative relationships between cytokine concentration and PMOS-like phenotypes from cohort 1. Splenocytes from the genetically diverse CTRL and LET cohorts were stimulated with anti-CD3/28 or LPS. Conditioned media was assayed for cytokines by multiplex analysis. Red denotes strong, positive correlation. White denotes no correlation. Blue denotes strong, negative correlation. Asterisk (*) denotes statistically significant correlations.

**Supplemental Figure 5.**
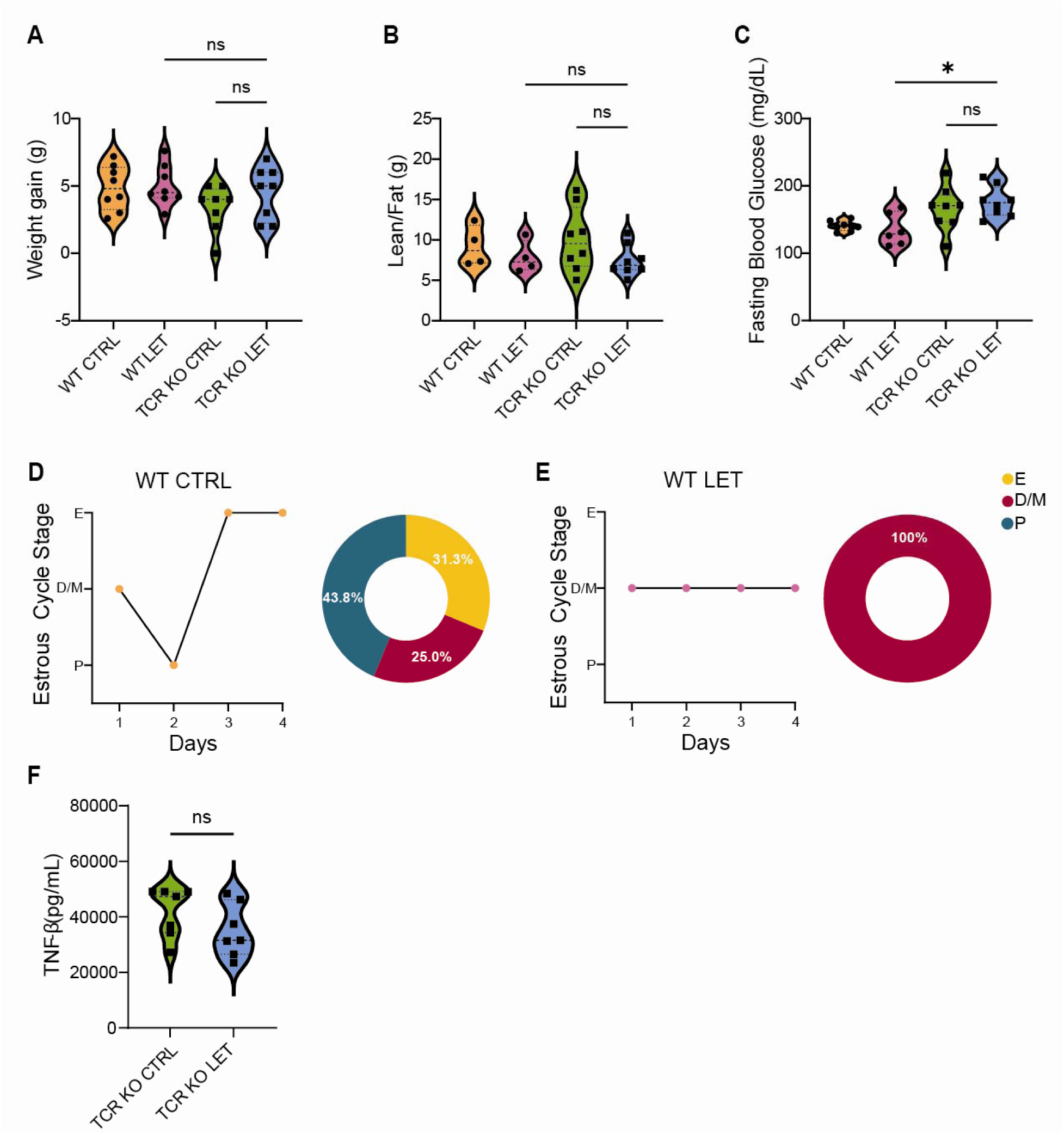
The adult paradigm of LET treatment does not result in a strong metabolic phenotype. (**A**) Total weight gain over the 5-week placebo or LET pellet treatment, (**B**) lean to fat mass ratio, and (**C**) fasting blood glucose from cohort 2. (**D-E**) Representative plots and percentage of mice cycling defined as progressing through all estrous cycle stages in order of WT CTRL (D) and WT LET (E). (**F**) Levels of TNFβ (pg/mL) in serum of TCR KO CTRL and LET cohorts.

**Supplemental Figure 6.**
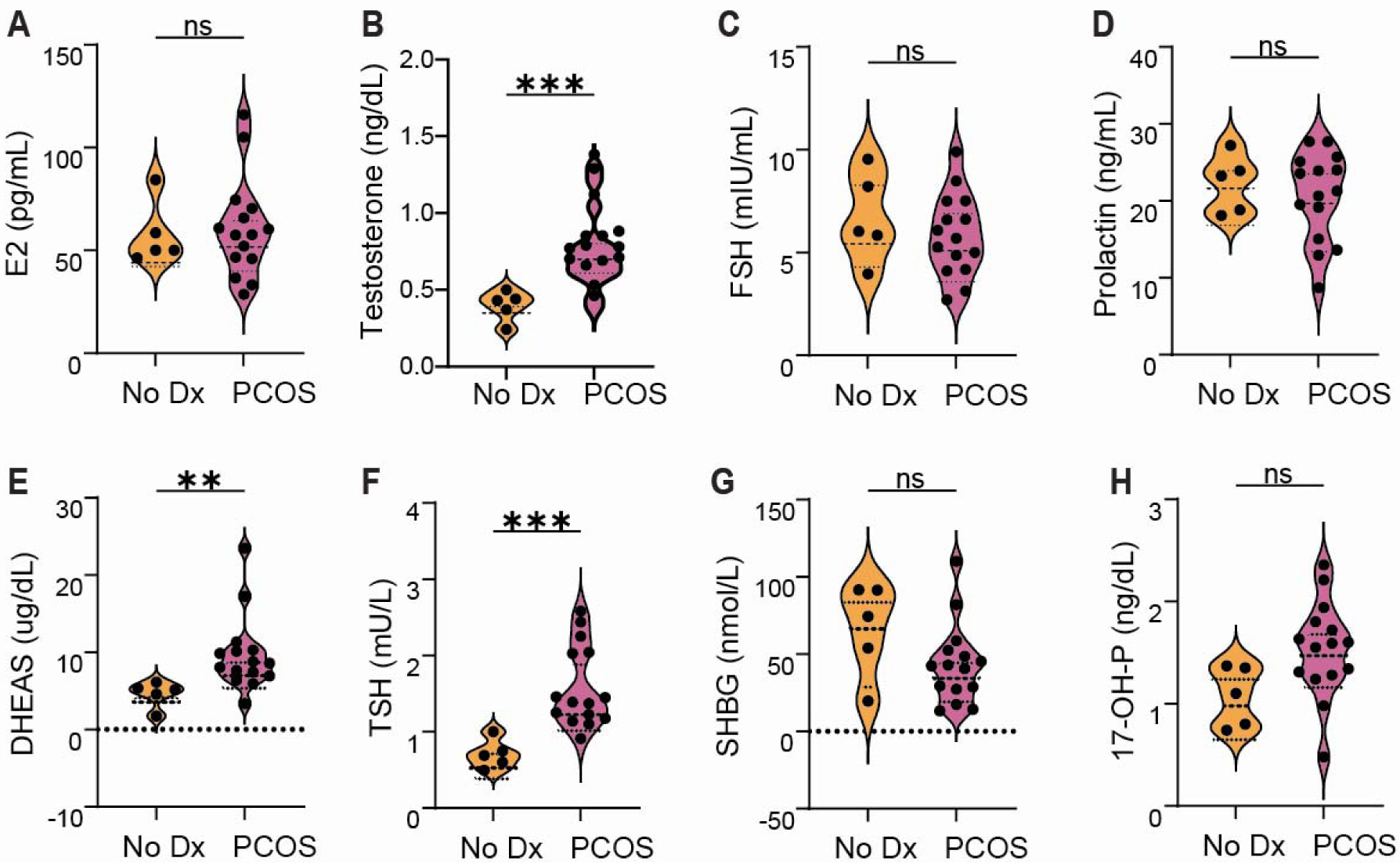
Hormone profiles of PMOS cohort are consistent with PMOS diagnosis. (**A-H**) Clinical parameters measured in women without PMOS (n = 5) or with PMOS (n = 15). Blood levels of (A) Estradiol (pg/mL), (B) Testosterone (ng/dL), (C) Follicular-stimulating hormone (mIU/mL), (D) Prolactin (ng/mL), (E) Dehydroepiandrosterone sulfate hormone (ug/dL), (F) Thyroid-stimulating hormone (mU/L), (G) Sex hormone-binding globulin (nmol/L) Cholesterol (mg/dL), and (H) 17-Hydroxyprogesterone (ng/dL). Data are presented as mean ± SEM and analyzed with t-test for parametric data or Mann-Whitney test for nonparametric data. Significance was accepted at p<0.05 and is indicated with an *.

**Supplemental Figure 7.**
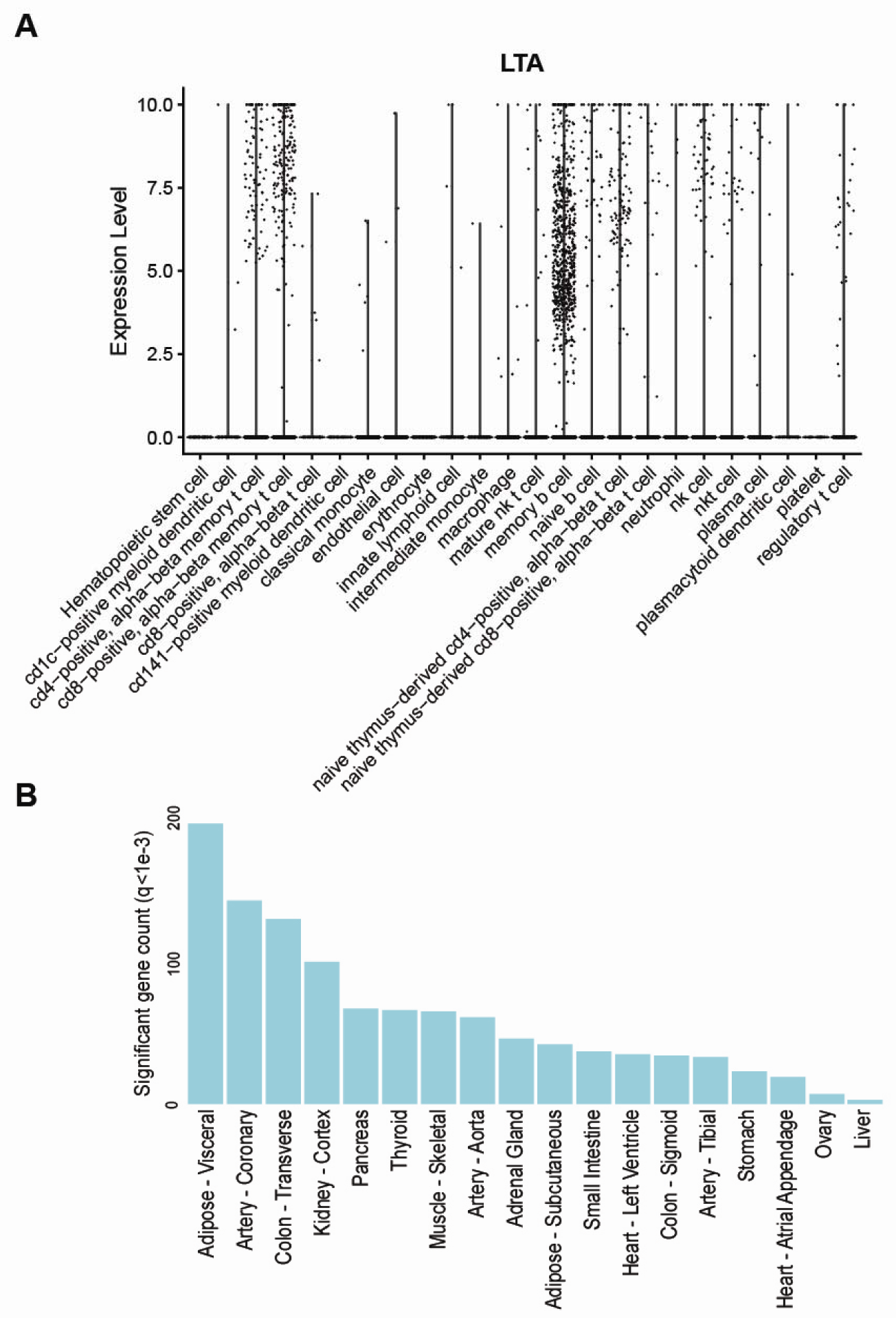
LTA expression in the spleen correlates with gene expression in the adipose. (**A**) Genotype-tissue expression levels of lymphotoxin-alpha (LTA), also known as TNF-β, from scRNAseq data of spleen from 31 premenopausal women (age <50) (**B**) Significant gene count of LTA in listed tissues from the same dataset.

**Supplemental Figure 8.**
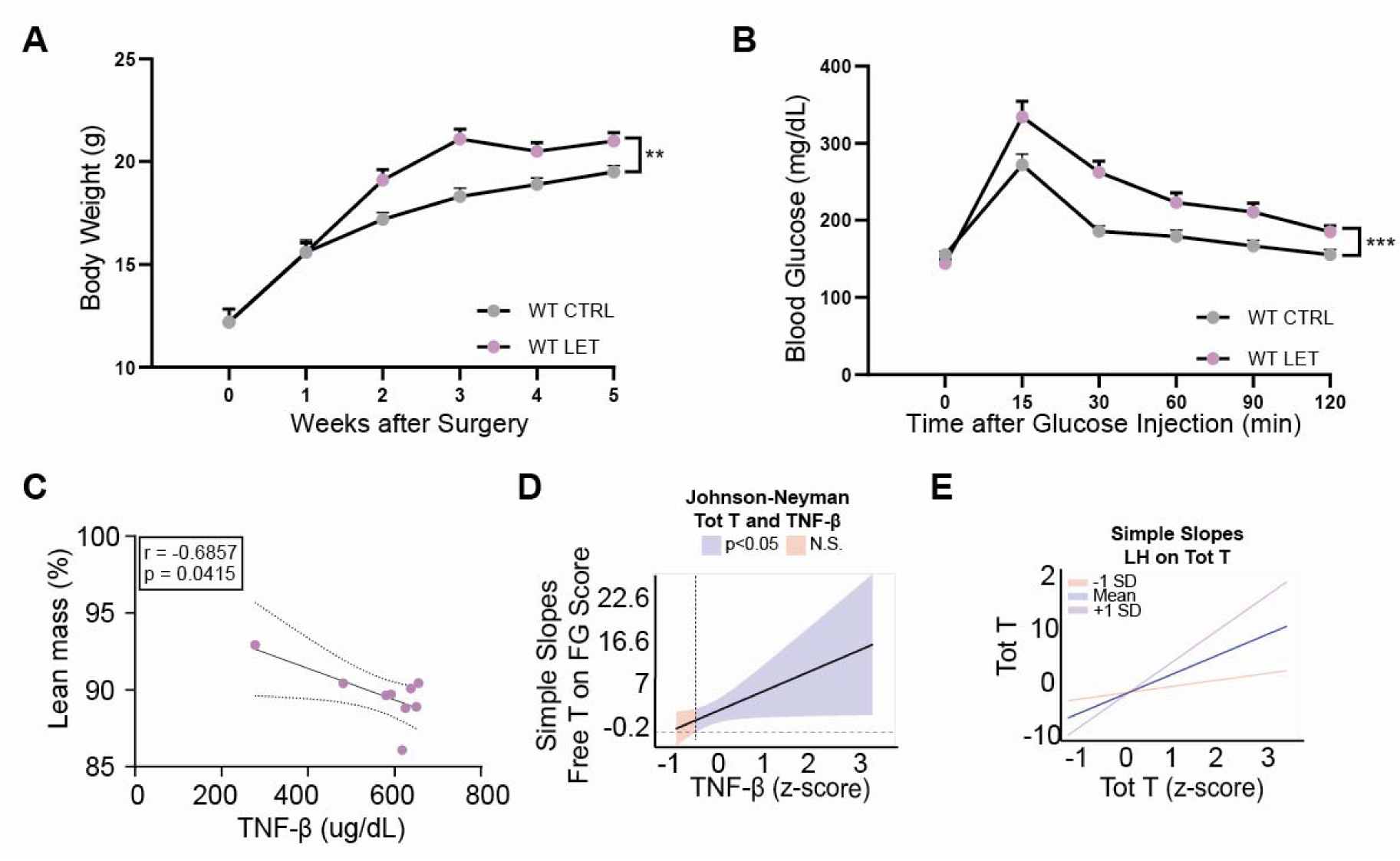
Pre-pubertal LET model has a metabolic phenotype. (**A**) Total weight gain of cohort 3 over the 5-week placebo or LET pellet treatment. (**B**) Blood glucose levels (mg/dL) over 120 minutes. (**C**) Correlations of TNFβ and lean mass (%) in WT LET cohort. (**D-E**) Moderation analysis of TNF-β on the relationship of free testosterone to Ferriman-Gallwey score.

**Supplemental Figure 9.**
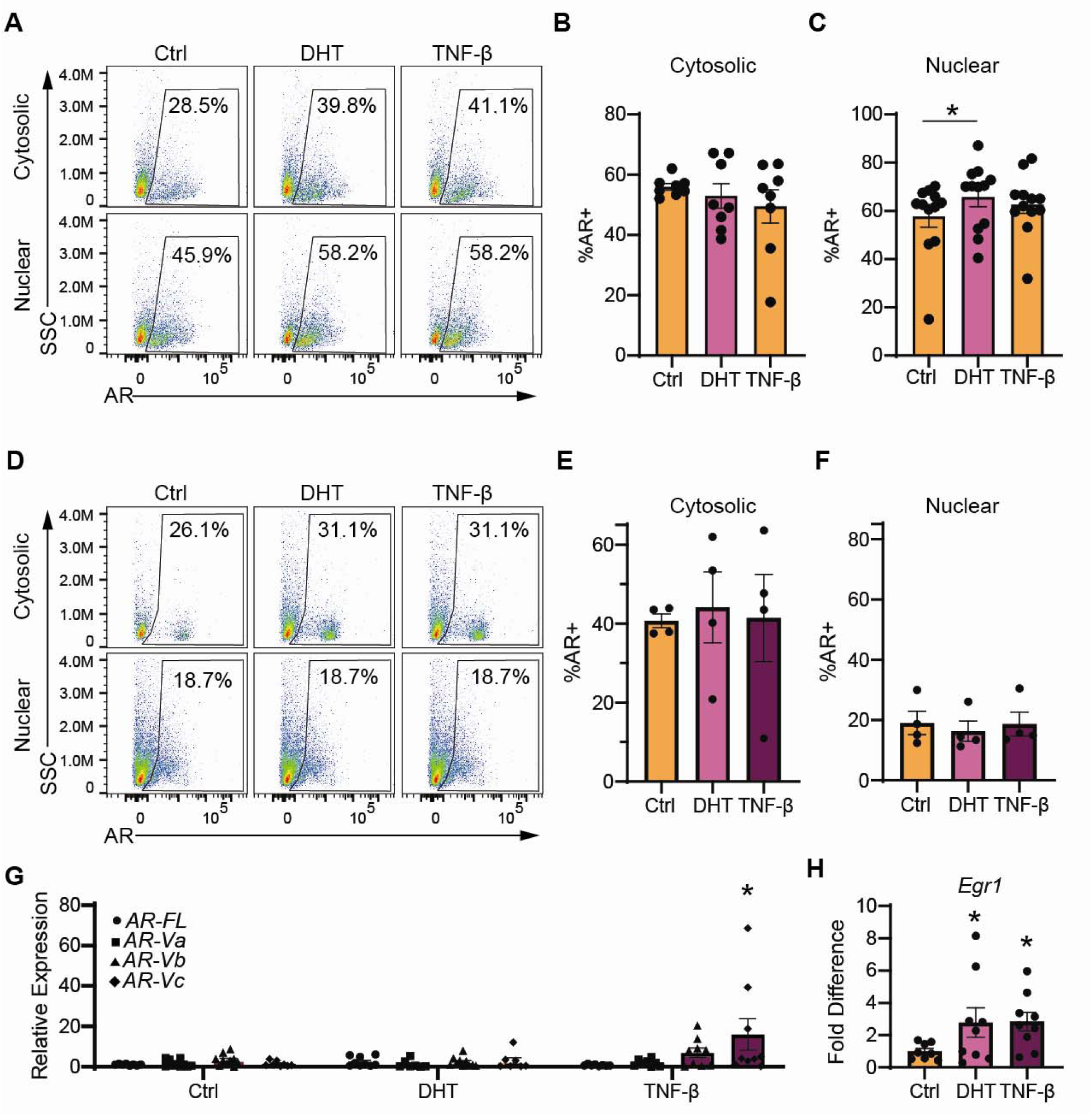
DHT activation of LβT2 cells confirms TNF-β effect on AR splice variants induction. (**A**) Representative flow cytometry gating of androgen receptor within nuclear and cytosolic components of LβT2 cells treated with DHT and TNF-β. (**B-C**) Quantification of flow cytometry in (A). (**D**) Representative flow cytometry gating of androgen receptor within nuclear and cytosolic compartments of mice ovarian cultures treated with DHT and TNF-β. (**E-F**) Quantification of flow cytometry in (D). (**G**) Relative expression of Ar splice variants in LβT2 cells treated with DHT and TNF-β. (**H**) Fold increase in mRNA of *Egr1* in LβT2 cells treated with DHT and TNF-β.

## Supplementary Tables

**Supplemental Table 1.** Summary of Mice used in Genetic LET-induce PMOS Screen.

| Mouse strain | Treatment | Cage N° | n Mice | Source |
| --- | --- | --- | --- | --- |
| A/J | ctrl | 39 | 4 | Jackson Labs |
| A/J | PMOS | 40 | 4 | Jackson Labs |
| BALB/cj | ctrl | 41 | 4 | Jackson Labs |
| BALB/cj | PMOS | 42 | 4 | Jackson Labs |
| CH3/Hej | ctrl | 43 | 4 | Jackson Labs |
| CH3/Hej | PMOS | 44 | 4 | Jackson Labs |
| DBA/2J | ctrl | 36 | 2 | Jackson Labs |
| DBA/2J | ctrl | 51 | 4 | Jackson Labs |
| DBA/2J | PMOS | 35 | 2 | Jackson Labs |
| DBA/2J | PMOS | 52 | 4 | Jackson Labs |
| NOD | ctrl | 37 | 4 | Jackson Labs |
| NOD | PMOS | 38 | 4 | Jackson Labs |
| BXD124 | ctrl | 34 | 4 | Williams, Ashbrook |
| BXD124 | PMOS | 17 | 3 | Williams, Ashbrook |
| BXD124 | PMOS | 33 | 4 | Williams, Ashbrook |
| BXD125 | ctrl | 27 | 2 | Williams, Ashbrook |
| BXD125 | PMOS | 26 | 3 | Williams, Ashbrook |
| BXD128 | ctrl | 1 | 2 | Williams, Ashbrook |
| BXD128a | ctrl | 21 | 3 | Williams, Ashbrook |
| BXD128a | PMOS | 20 | 3 | Williams, Ashbrook |
| BXD44 | PMOS | 6 | 2 | Williams, Ashbrook |
| BXD48a | ctrl | 23 | 2 | Williams, Ashbrook |
| BXD48a | PMOS | 22 | 2 | Williams, Ashbrook |
| BXD51 | ctrl | 25 | 2 | Williams, Ashbrook |
| BXD51 | PMOS | 24 | 2 | Williams, Ashbrook |
| BXD60 | ctrl | 9 | 2 | Williams, Ashbrook |
| BXD60 | ctrl | 16 | 3 | Williams, Ashbrook |
| BXD60 | PMOS | 8 | 3 | Williams, Ashbrook |
| BXD70 | PMOS | 14 | 3 | Williams, Ashbrook |
| BXD71 | ctrl | 4 | 3 | Williams, Ashbrook |
| BXD73 | ctrl | 11 | 2 | Williams, Ashbrook |
| BXD73 | PMOS | 10 | 2 | Williams, Ashbrook |
| BXD73b | ctrl | 30 | 3 | Williams, Ashbrook |
| BXD73b | PMOS | 29 | 4 | Williams, Ashbrook |
| BXD75 | ctrl | 5 | 2 | Williams, Ashbrook |
| BXD75 | PMOS | 3 | 2 | Williams, Ashbrook |
| BXD77 | ctrl | 28 | 3 | Williams, Ashbrook |
| BXD77 | PMOS | 7 | 3 | Williams, Ashbrook |
| BXD78 | ctrl | 2 | 2 | Williams, Ashbrook |
| BXD79 | ctrl | 19 | 2 | Williams, Ashbrook |
| BXD79 | PMOS | 18 | 2 | Williams, Ashbrook |
| BXD89 | ctrl | 13 | 2 | Williams, Ashbrook |
| BXD89 | PMOS | 12 | 2 | Williams, Ashbrook |
| BXD89 | PMOS | 15 | 3 | Williams, Ashbrook |

**Supplemental Table 2.** 18-Color Mouse Flow Cytometry Panel.

| Specificity | Fluorophore | Dilution | Purpose | Clone | Manufacturer | Catalog # |
| --- | --- | --- | --- | --- | --- | --- |
| Amine groups | Zombie NIR | 1:3200 | Viability | --- | Biolegend | 423106 |
| CD45 | BV 510 | 1:500 | Pan leukocyte | 30-F11 | Biolegend | 103138 |
| CD11b | PerCP-eFluor 710 | 1:400 | Myeloid cells | M1/70 | ThermoFisher | 46-0112-82 |
| Ly6C | PE-Cy7 | 1:1000 | Monocytes | HK1.4 | Biolegend | 128018 |
| NK1.1 | APC-FIRE 810 | 1:400 | NK cells | S17016D | Biolegend | 156519 |
| CD11c | Pacific Blue | 1:200 | Dendritic cells | N418 | Biolegend | 117322 |
| F4/80 | AF 488 | 1:1000 | Macrophages | BM8 | Biolegend | 123120 |
| CD19 | AF 647 | 1:1000 | B cells | 6D5 | Biolegend | 115522 |
| CD3 | PerCP | 1:50 | T cells | 145-2C11 | Biolegend | 100326 |
| CD4 | Nova Fluor Blue 610 | 1:200 | T helper cells | GK1.5 | ThermoFisher | M001T02B06-4 |
| CD8 | BV570 | 1:100 | T cytotoxic cells | 53-6.7 | Biolegend | 100740 |
| PD1/CD279 | BV 421 | 1:800 | Activated T cells | 29F.1A12 | Biolegend | 135218 |
| CD25 | BV605 | 1:200 | Activated Tregs | PC61 | Biolegend | 102036 |
| CD127 | PE | 1:400 | lo on Treg | S18006K | Biolegend | 158204 |
| CD62L | BV785 | 1:100 | Naive T cells | MEL-14 | Biolegend | 104440 |
| CD44 | AF700 | 1:50 | Memory T/B cells | IM7 | Biolegend | 103026 |
| MHCII | BV711 | 1:1000 | Activated B cells | M5/114.15.2 | Biolegend | 107643 |
| CD38 | BV650 | 1:1000 | Activated B cells | 90/CD38 | BD Biosciences | 740489 |

**Supplemental Table 3.** AR translocation Mouse Flow Cytometry Antibodies.

| Specificity | Fluorophore | Dilution | Purpose | Clone | Manufacturer | Catalog # |
| --- | --- | --- | --- | --- | --- | --- |
| Amine groups | Zombie NIR | 1:3200 | Viability | --- | Biolegend | 423106 |
| CD45 | AF 405 | 1:1000 | Pan leukocyte | 30-F11 | Biolegend | 103138 |
| AR | FITC | 1:500 | Androgen receptor | Polyclonal | ThermoFisher | ANDR-FITC |

## Notes

### Competing Interest Statement

The authors have declared no competing interest.

### Summary of Updates

This manuscript has been substantially revised with new data added and reorganization of figures and text.

